# RASSF effectors couple diverse RAS subfamily GTPases to the Hippo pathway

**DOI:** 10.1101/2020.02.05.923433

**Authors:** Dhanaraman Thillaivillalan, Swati Singh, Ryan C. Killoran, Anamika Singh, Xingjian Xu, Julia Shifman, Matthew J. Smith

## Abstract

Activated RAS GTPases signal by directly binding effector proteins. Effectors have a folded RAS association (RA) domain that binds exclusively to GTP-loaded RAS, but the specificity of most RA domains for >150 RAS superfamily GTPases is unknown. Ten RAS-association domain family (RASSF) proteins comprise the largest group of effectors, proposed to couple RAS to the pro-apoptotic Hippo pathway. We show that RASSF1-6 complex with Hippo kinase, while RASSF7-10 are a separate family related to p53-regulatory ASPP effectors. Only RASSF5 directly binds activated HRAS and KRAS. Structural modelling reveals that expansion of RASSFs in vertebrates included amino acid substitutions that alter their GTPase binding specificity. We demonstrate that the tumour suppressor RASSF1A complexes with the GTPases GEM, REM1, REM2 and the enigmatic RASL12. Interplay between RASSFs and RAS GTPases can drastically restrict YAP1 nuclear localization. Thus, these simple scaffolds can link activation of diverse RAS proteins to Hippo or p53 regulation.

## Introduction

RAS small GTPases are archetypal signalling proteins that control the direction and intensity of signals by acting as ‘molecular switches’. The proteins present distinct conformations dependent on a bound nucleotide: when bound to GTP they are ‘on’ and interact with diverse effector proteins that relay signals downstream. When GDP-bound, they lose their ability to interact with effectors and are considered ‘off’. Three RAS proteins (KRAS, NRAS and HRAS) have been intensely studied since the 1980s as key drivers of human cancers (1). They are members of a GTPase superfamily that consists of >150 proteins divided into the RAS, RHO, ARF, RAB, and RAN branches (2). The RAS subfamily consists of 35 small GTPases in humans, yet the function of most remains understudied. They can be ostensibly classified into two groups: those that promote cell proliferation or survival (RAS, RAL (3), RIT (4), RHEB (5) and RAP (6)) and those that induce cell cycle arrest or apoptosis (RASD (7), NKIRAS (8), RASL (9), DIRAS (10) and RGK (11)). Comparatively little is known about how these later tumour suppressor RAS GTPases function.

Effector proteins determine where activated GTPase signals are routed, and RAS-induced tumourigenesis is known to depend on multiple effectors (12, 13). Effectors compete for activated RAS using recognition domains called <u>R</u>AS-<u>A</u>ssociation (RA) or <u>R</u>AS <u>B</u>inding <u>D</u>omains (RBDs) (collectively referred to here as RA domains). There are >50 RA domains in the human proteome by sequence homology (14), and all proteins comprising these domains are currently deemed ‘RAS effectors’. Despite this, there is likely extensive plasticity in effector-GTPase interactions whereby each effector signals downstream of multiple GTPases. This idea is supported by data showing PI3K binds to multiple distinct RAS subfamily GTPases (15).

Paradoxically, H/K/N-RAS binding to the poorly studied <u>R</u>AS <u>ass</u>ociation domain family (RASSF) of effectors is suggested to promote apoptosis rather than proliferation (16) and the promoters of ten genes encoding RASSF proteins are hypermethylated in numerous cancers (17, 18). These are amongst the most frequently inactivated tumour suppressors, candidate biomarkers and appealing therapeutic targets. With ten homologous family members, they also comprise one-fifth of the proposed RAS effector landscape. Co-expression of RASSF1 or RASSF5 with oncogenic RAS triggers an apoptotic response (19, 20), and several RASSF members inhibit proliferation of tumour-derived cells (21–24). Consistent with a tumour suppressor role, mice lacking RASSF1 are prone to spontaneous tumorigenesis (25). The proteins are simple scaffolds and can be divided into two groups: C-terminal or classical (C-)RASSFs 1-6 and N-terminal (N-)RASSFs 7-10 (**Figure 1A**). All family members have an RA domain proposed to bind H/K/N-RAS. Some evidence for this exists for RASSF1 (20), RASSF2 (26), RASSF4 (22), RASSF5 (16) and RASSF7 (27), but only the RASSF5 interaction is supported by *in vitro* binding data and a structure of its RA domain bound to HRAS (28). Other studies have linked RASSFs to the small GTPases RAP1A (29), RHEB (30), RHOA (31), and RAN (32). Conclusive biophysical evidence for these interactions is lacking. Besides an RA domain, C-RASSFs comprise a helical SARAH motif that directly binds the pro-apoptotic Hippo kinases MST1/2 (33–36), which direct the cellular response contact inhibition, mechanical tension and polarity to control organ size and tissue homeostasis *via* coordinated apoptosis. Though RASSFs are one of few direct Hippo interaction partners, their impact on MST1/2 activity or YAP/TAZ nuclear translocation has not been fully investigated. The N-RASSFs lack a SARAH motif and whether they have a role in Hippo signalling is unclear.

**Figure 1.**
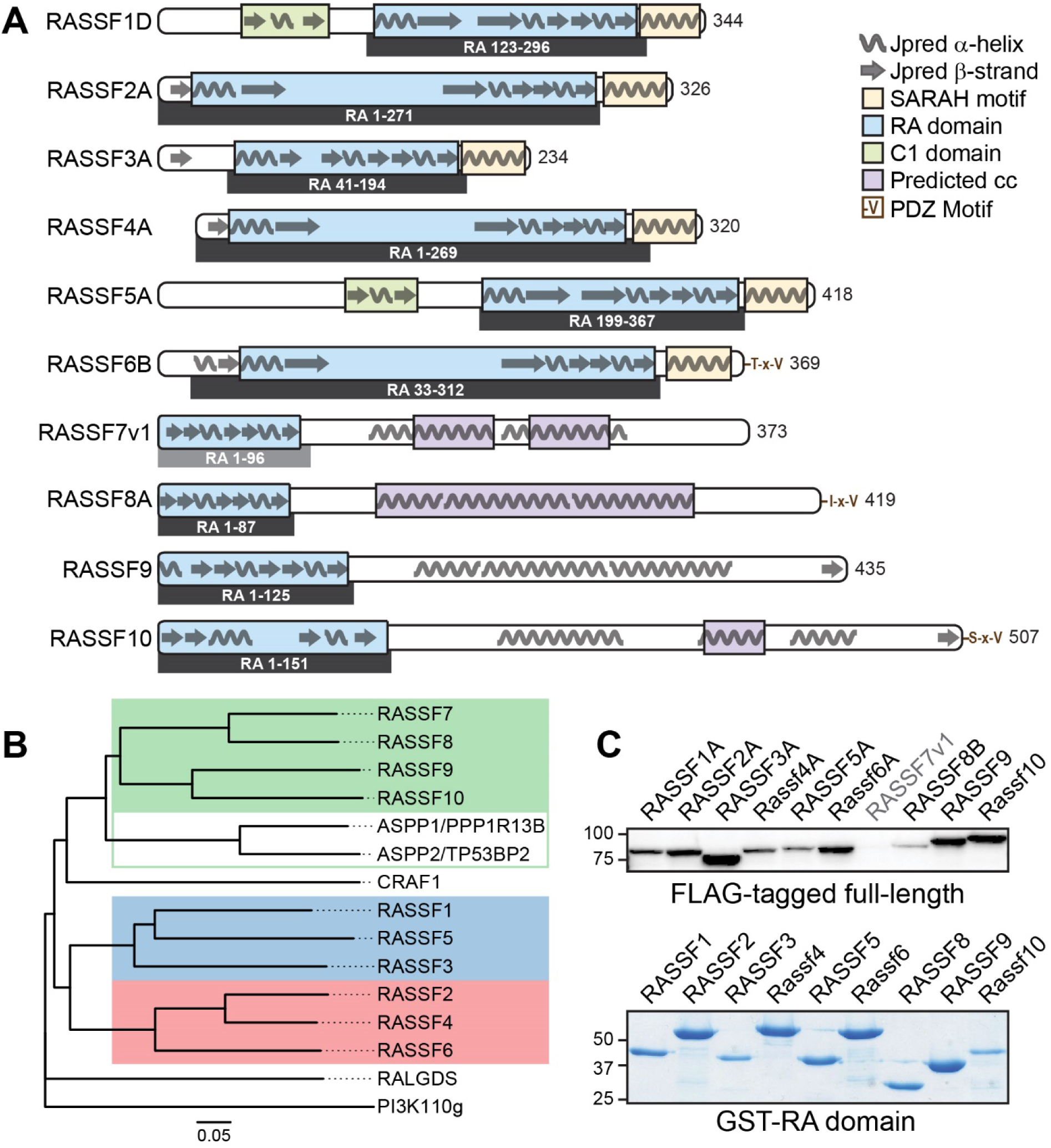
Domain architecture and purification of RASSF effectors. **(A)** Schematic representation of the RASSF family members. The longest isoform of each RASSF is depicted, overlaid with secondary structure predictions (Jpred). RA domains are in blue, and amino acid positions of the domain expression constructs are detailed below. C-RASSFs have helical SARAH (Salvador-Rassf-Hippo) motifs, in yellow. RASSF1 and RASSF5 have C1 zinc finger domains, in green. SMART-predicted coiled coil domains of the N-RASSFs are in purple. **(B)** Classification of RASSF homologs based on amino acid sequence similarity. RA domains of the C- and N-RASSFs were aligned with RA domains from ASPP1, ASPP2, CRAF-1, RALGDS and PI3Kγ. The N-RASSF family (green) clusters with the two ASPP domains. Two clusters are evident in the C-RASSF family, one composed of RASSF1/3/5 (blue) and another of RASSF2/4/6 (red). **(C)** anti-FLAG Western blot shows expression of full-length, FLAG-tagged RASSF isoforms (top) and a Coomassie-stained SDS-PAGE gel shows the purified, GST-tagged RA domain proteins (bottom).

A family of tumour suppressors directly linking RAS with apoptotic Hippo signalling is highly intriguing and requires definitive analysis. More generally, the specificity of effector RA domains for RAS or alternative RAS-like GTPases remains unknown. Here, we begin to resolve these questions for the RASSF family of purported RAS effectors. We clarify which RASSF effectors directly complex with activated KRAS and define interaction partners for the associated functional domains in these scaffolds. New RAS subfamily interaction partners are identified for the important RASSF1A tumour suppressor that support a role in cell cycle regulation and calcium (Ca^2+^) homeostasis. We reveal that RASSFs do not inherently synergize with oncogenic RAS to promote apoptosis, and connect these new data to Hippo activation.

## Results

### Delineation and Purification of RASSF RA domains

To investigate RASSF interactions with GTPases required purified and well-folded RA domains. Only the RASSF5 RA domain has been previously characterised. To define domain boundaries for the other nine RASSFs we used a combination of sequence homology, secondary structure predictions, and the understanding that effector RAS-binding domains share a ubiquitin superfold (ββαββαβ). Additionally, the RA domain of RASSF5 has a distinct N-terminal α-helix that is essential for binding RAS (28). **Figure 1A** shows the domain architecture of the longest isoform of each RASSF and the boundaries of their RA, C1 and SARAH domains overlaid with secondary structure predictions (JPred). By homology, RA domains from the N-RASSFs (7-10) do not cluster with C-RASSFs, but rather with RA domains from two <u>a</u>poptosis-<u>s</u>timulating <u>p</u>roteins of <u>p</u>53 (ASPP) effectors (also PPP1R13B and TP53BP2; **Figures 1B and S1A**) (37, 38). Further, N-RASSFs have a long coiled-coil region analogous to those found in ASPP proteins (**Figure S1B**) and do not encode the αN-helix present in RASSF5. We were able to express and purify RA domains from RASSFs 8-10 as recombinant GST-fusion proteins, but the RASSF7 domain proved insoluble under numerous expression and purification conditions and was not considered further (**Figure S2**). Amongst the C-terminal RASSF proteins, RASSF1, RASSF3, and RASSF5 form one homology subgroup and RASSF2, RASSF4, and RASSF6 a second. Each of the RASSF1/3/5 RA domains encode an αN-helix critical to RAS binding, and production of recombinant protein using our defined boundaries resulted in RA domains that could be purified to high concentration and homogeneity (**Figure S2**). The RASSF2/4/6 subgroup each show a region of predicted secondary structure at their N-termini that bears no homology to RASSF C1 domains. In fact, the sequences encoding these elements are homologous to the αN-helix and β1 strands of RASSF1/3/5 (**Figure S3**). Indeed, RASSF2/4/6 RA domains lacking these N-terminal regions proved highly insoluble, but their inclusion stabilized recombinant proteins (**Figure S2**). Thus, it appears that long unstructured segments have evolved between strands β1 and β2 of the RASSF2/4/6 RA domains with unknown consequences on protein folding. The analogous region of the *D. melanogaster* ortholog is separated by over 500 amino acids (**Figure S3**). Delineation of these domain boundaries and purification of recombinant RA domains from nine RASSF homologs provided an opportunity to systematically assess their ability to bind RAS GTPases. We also generated vectors to express full-length, FLAG-tagged RASSF proteins in cultured cells, focusing on the longest isoform for each (**Figure 1C**). As with its RA domain we found full-length RASSF7 mostly insoluble, but now had a toolbox to assay interactions for the other nine RASSF effectors and sought to identify binding partners for all functional regions.

### Differential RASSF Interactions with KRAS, Hippo Kinase and ASPP Effectors

RASSF effectors are small scaffolds with few recognized domains outside the RA module. An exception is the SARAH motif at the C-termini of all C-RASSFs (**Figure 1A**). Previous studies revealed interactions between the SARAH motifs of RASSF1, RASSF2 and RASSF5 with those of the mammalian Hippo kinases (34, 39, 40). To ascertain if all RASSF proteins interact with the Hippo kinase MST1 we co-expressed full-length FLAG-RASSF proteins with GFP-tagged MST1 and performed co-immunoprecipitations (co-IPs). This approach established that all six C-RASSF proteins (RASSF1-6) associate with MST1 (**Figure 2A**). Further, we consistently observed increased C-RASSF protein levels upon co-expression with MST1, suggesting the kinase promotes RASSF stability. A RASSF5 construct truncated after the RA domain confirmed that loss of the SARAH motif disrupts MST1 binding. The N-RASSFs 8-10 lack a defined SARAH motif and did not co-precipitate with the Hippo kinase. These data reveal that all C-RASSFs can complex with MST1 and may regulate Hippo signalling.

**Figure 2.**
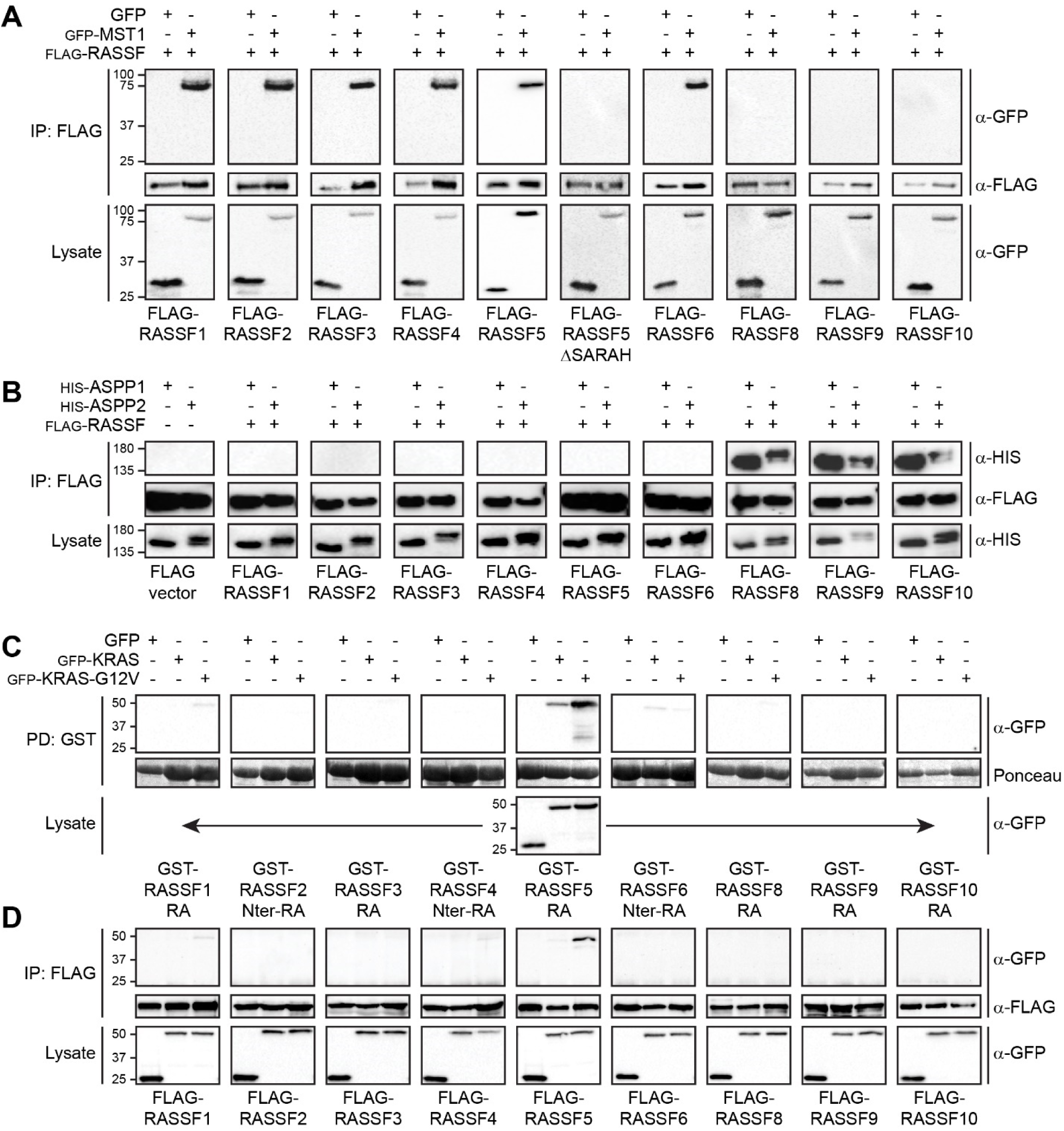
Differential interactions between RASSF effectors and KRAS, MST1 kinase and the ASPP effectors. **(A)** Complex of MST1 kinase and RASSF effectors. Cells were co-transfected with GFP alone or GFP-MST1 and FLAG-RASSF proteins. Anti-FLAG antibodies were used to immunoprecipitate RASSFs and interacting proteins were revealed by anti-GFP immunoblot (top). FLAG-RASSF5 ΔSARAH demonstrates dependency on the SARAH motif. **(B)** Interactions between ASPP1/2 effectors and the RASSF proteins. Cells were co-transfected with HIS-tagged ASPP1 or ASPP2 with FLAG-RASSF proteins. Anti-FLAG immunoprecipitations were probed by anti-HIS immunoblot to reveal interactions (top). **(C)** Binding of purified RASSF RA domains to KRAS. Cells expressing GFP alone, GFP-KRAS-WT or GFP-G12V-KRAS were lysed and incubated with purified GST-RA domains. anti-GFP immunoblots (top) revealed bound proteins. A ponceau stain shows GST-RA domains. **(D)** Interaction between KRAS and full-length RASSF proteins. Cells were co-transfected with GFP alone, GFP-WT-KRAS or GFP-G12V-KRAS with FLAG-RASSF proteins. Anti-FLAG antibodies were used to precipitate RASSFs, and anti-GFP immunoblots reveal interacting proteins (top).

As N-RASSFs exhibit greater homology to the ASPP family of purported RAS effectors, and these are capable of oligomerization (41), we investigated whether RASSFs 8-10 could complex with ASPP1/2. Indeed, the *Drosophila* ortholog dRASSF8 associates with dASPP to regulate cell adhesion (42). We co-expressed nine FLAG-RASSF proteins with either HIS-tagged ASPP1 or ASPP2 and performed co-IPs. All three N-RASSFs co-precipitated with both ASPP1 and ASPP2, while none of the C-RASSFs proteins bound (**Figure 2B**). Interestingly, the ASPP effectors are direct regulators of p53 and known tumor suppressors with a pro-apoptotic function, not unlike RASSFs.

While numerous studies have probed individual interactions between RASSF family members and RAS GTPases, results have been ambiguous and have not employed isolated RA domains. Further, our sequence alignment reveals only RASSF1 has a conserved Lys residue required for RASSF5 binding to HRAS (Lys288; **Figure S3**). To resolve whether all RASSF homologs are direct RAS effectors we systematically probed interactions between activated KRAS and the nine purified RA domains. GST-tagged RA domains were added to cell lysates expressing full-length, GFP-tagged WT-KRAS (wild-type), G12V-KRAS (constitutively activated) or GFP alone (control) and complexes isolated on glutathione beads. For a proper comparative analysis, each RA domain was purified by size exclusion chromatography and added to the same batch of cell lysate at a final concentration of 1 μM. GST pull-downs were washed 5X in the identical buffer, and anti-GFP immunoblots exposed in parallel. As expected, we detected strong binding between the RASSF5 RA domain and G12V-KRAS (**Figure 2C**), but no other RASSF complexed with the GTPase. As this approach only assayed interactions with our defined RA domains, we repeated this analysis using the longest isoform of each RASSF. We performed anti-FLAG precipitations following co-expression of FLAG-RASSF proteins with GFP-tagged WT-KRAS, G12V-KRAS or GFP alone. As with the GST binding assays, only RASSF5 interacted with G12V-KRAS (**Figure 2D**). We consistently observed weak association between RASSF1 and G12V-KRAS, and to a lesser extent RASSF4 and RASSF6, but this was only evident upon long exposure of immunoblots. That only RASSF5 is able to bind KRAS contradicts much of the literature, but is consistent with some data suggesting other RASSFs have reduced affinity for RAS (43).

### RAS-Induced Apoptosis and RASSF Interactions with RAS Superfamily Small GTPases

The lack of KRAS binding suggested that all RASSF effectors may not elicit an apoptotic response downstream of RAS *via* a direct mechanism, as previously proposed. To test this, we systematically assessed the capacity for RASSF proteins to stimulate cell death. We measured apoptosis by Annexin V staining of HEK 293T cells expressing FLAG-tagged RASSF proteins alone or together with G12V-KRAS. FACS analysis showed that no RASSF proteins intrinsically stimulate apoptosis over control levels (**Figure 3A/B**). Moreover, we observed no significant increase in Annexin V staining when RASSFs were co-expressed with activated KRAS, which alone stimulated a 4-fold increase in apoptosis (50-60% of KRAS-expressing cells were apoptotic). The homologous HRAS protein (G12V) also induced a significant 3.6-fold increase in apoptosis, but this was not augmented at all by co-expression of RASSF5. These results contradict a model whereby RASSF proteins synergize with oncogenic RAS to induce cell death, and validate that these effectors are not universal RAS-binders.

**Figure 3.**
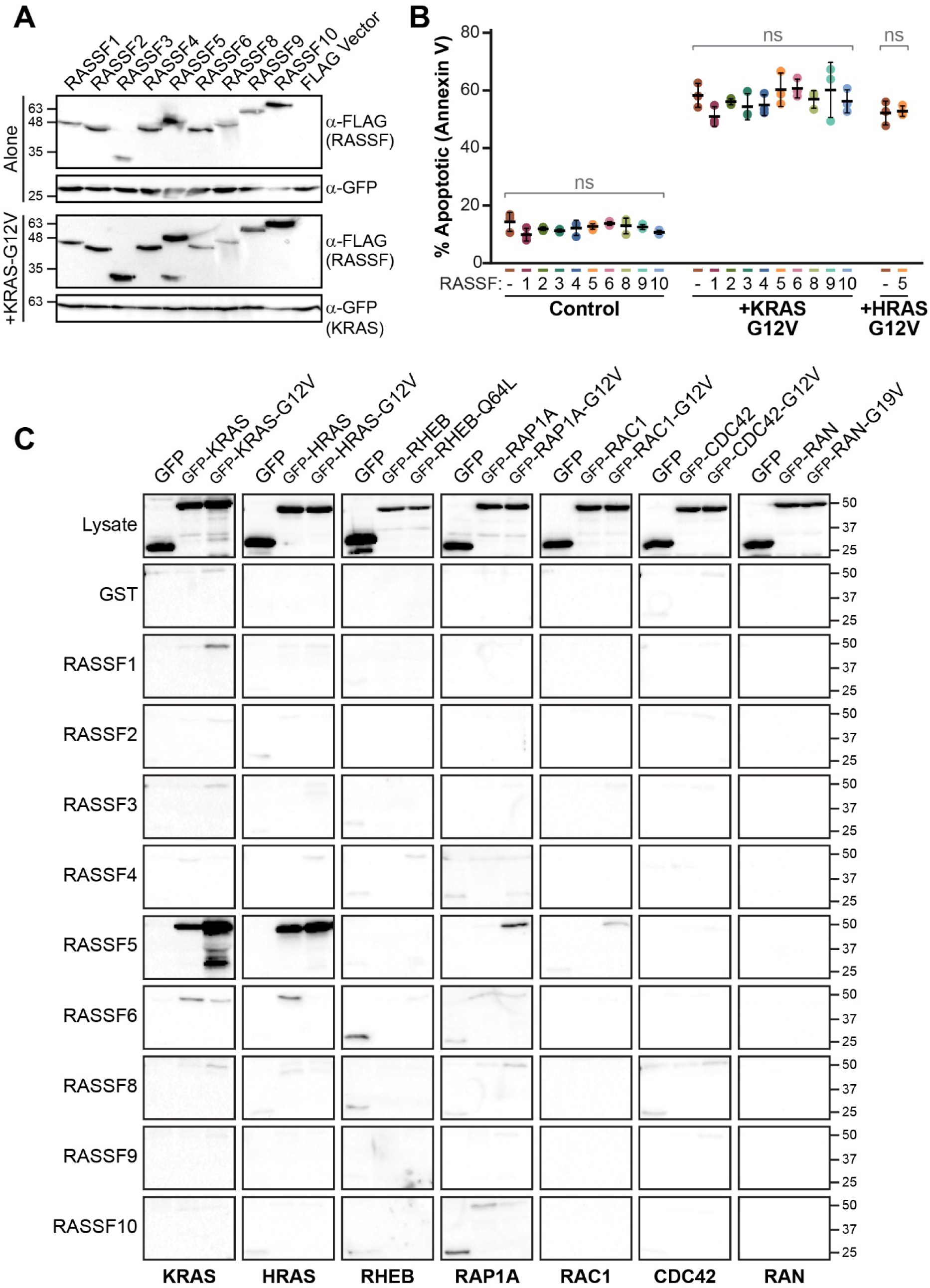
KRAS-induced apoptosis and RASSF RA domain binding to candidate GTPases. **(A)** HEK 293T cells expressing full-length FLAG-RASSF proteins alone or with GFP-tagged, activated KRAS or HRAS were assessed for apoptosis using Annexin V staining. A fraction of cells assayed for apoptotic activity verified expression levels of FLAG-tagged RASSF and GFP-tagged GTPase proteins. Immunoblots with anti-FLAG and anti-GFP are shown**. (B)** RASSFs do not intrinsically stimulate apoptosis. RASSFs did not significantly induce apoptosis over controls, nor did they augment the apoptotic response observed in G12V-KRAS or G12V-HRAS expressing cells (ns=not significant as measured by two-way ANOVA). **(C)** Probed interactions between the candidate GTPases and RA domains from nine RASSF homologs. Lysates expressing GFP alone or GFP-tagged wild type/activated mutants of the GTPases KRAS (control), HRAS, RHEB, RAP1A, RAC1, CDC42, or RAN were incubated with purified GST-RA domains. Complexes on glutathione beads were separated by SDS-PAGE and analyzed by anti-GFP immunoblot.

We presumed that RASSF effectors (other than RASSF5) might associate with distinct members of the RAS superfamily. Using discovery approaches to elucidate GTPase interactions is difficult as the proteins must be highly expressed, be in the active/GTP-bound conformation and not engaged with other high affinity binders. Thus, we took a systematic approach by first selecting six candidate GTPases to perform comparative binding assays: HRAS, RAP1A, RHEB, RAN (all previously proposed to bind RASSFs (29, 30, 32)), and the RHO family GTPases RAC1 and CDC42 (as overexpression of RASSFs alters cell morphology). HRAS is identical to KRAS across its two RA domain binding sites (switch I and II), while the five other GTPases have amino acid substitutions at several key residues. To examine binding to RASSFs we generated expression constructs for wild-type or activated mutant (based on RAS G12V) variants. Purified, GST-tagged RA domains from the nine RASSFs were added to cell lysates expressing these GTPases, and complexes isolated on glutathione beads. As expected, RASSF5 bound to KRAS and HRAS, and weakly to the homolog RAP1A, but our results did not corroborate a single interaction between other family members and any GTPase (**Figure 3C**). Thus, though RASSFs comprise an RA domain our data suggest upstream GTPase signals feeding into these tumour suppressors, and the cellular manifestation of these signals, are largely unknown.

### RASSF1 is an Effector of the RGK Family of RAS GTPases

To develop a second set of candidates we considered how residues in RASSF RA domains complement those in small GTPases. RAS effectors share a conserved mode of RAS recognition based on formation of an intermolecular, anti-parallel β-sheet (between β2 and β3 of RAS and β1, β2 and α1 of the RA domain), and RASSF5 has a distinctive αN-helix that contacts the RAS switch II region. Having established positions of secondary structure for all RASSF RA domains (**Figure S3**) we used the crystal structure of RASSF5-HRAS (**Figure 4A**) to construct structural models for the six C-RASSF RA domains (**Figures 4B** and **S4**). Models for the N-RASSF domains were generated based on an AFDN-HRAS structure (14). Residues in each αN, β1 and β2 region specify small GTPase interactions based on archetypal RAS-effector binding. Here, we focus on identifying novel GTPase interactors for RASSF1, as it has become an intriguing clinical biomarker and retains most of the key RAS-binding residues from its paralog, RASSF5. The only significant amino acid substitution in RASSF1 is an Asn in the αN-helix. This is a Cys in RASSF5 that mediates a key hydrophobic interaction with HRAS (**Figure 4A/B**). To explore the importance of this residue as a specificity determinant, we performed *in vitro* binding experiments using isothermal titration calorimetry (ITC). We measured a dissociation constant (*K*_d_) for wild-type RASSF5 RA domain binding to GMPPNP-loaded KRAS of 1.7 μM. A C225N mutant of RASSF5, mimicking RASSF1, had a 7-fold weaker *K*_d_ of 11.3 μM (**Figure 4C**). Conversely, wild-type RASSF1 RA domain had no measurable affinity for KRAS but a swapped N149C αN mutation enabled this interaction, with a *K*_d_ of 9.6 μM (**Figure 4D**). Taken together, amino acid substitutions in RASSF αN-helices significantly impact binding to small GTPases and, along with residues in β1 and β2, are key specificity determinants.

**Figure 4.**
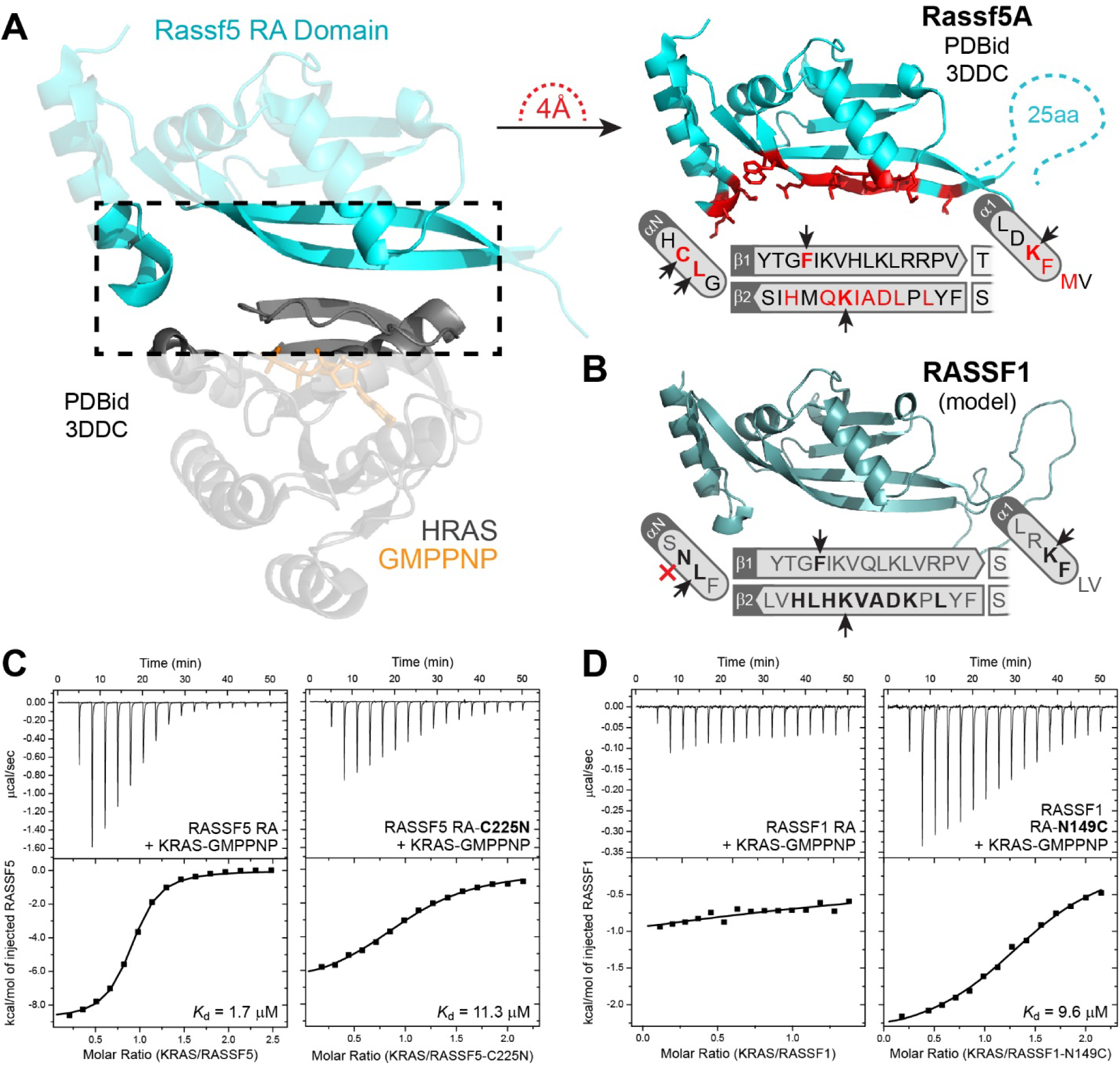
Structural modelling of RASSF RA domains elucidates specificity-determining substitutions in the αN-helix. **(A)** Ribbons diagram of the HRAS (grey)-RASSF5 (blue) structure (PDBid 3DDC), highlighting the interface (left). A ribbons diagram of the RASSF5 RA domain and a schematic showing amino acid positions in the β1 and β2 strands, and the αN and α1 helices is at right. Residues within 4 Å of HRAS are highlighted in red. Amino acids indicated by a black arrow are predicted to have side chains that interact with RAS. **(B)** Structural model of the RASSF1 RA domain. Homology modeling based on the HRAS-RASSF5 structure shows the high similarity between RASSF1 and RASSF5 at the GTPase binding interface. RASSF1 residues predicted within 4 Å of HRAS are in bold. Black arrows indicate key conserved amino acids with RASSF5, while a red ‘X’ denotes a Cys-to-Asn substitution in αN. (**C)** Binding thermodynamics of the RASSF-KRAS interaction. ITC determined the RASSF5 RA domain interaction with KRAS-GMPPNP is 1.7 μM (left). A Cys-to-Asn mutation in RASSF5 αN weakens the interaction 7-fold (right). **(D)** ITC shows the RASSF1 RA domain does not measurably interact with KRAS-GMPPNP (left), but a RASSF5-mimetick Asn-to-Cys αN mutant interacts with a *K*_d_ of 9.6 μM (right).

To identify small GTPase partners for RASSF1 we considered the three contact sites that define a RASSF-GTPase complex. As the RASSF1 β1, β2 and α1 regions are nearly identical to RASSF5 (**Figure 5A**: contact sites 2 and 3), we focused on the αN-helix Cys-Asn substitution (contact site 1). We aligned the amino acid sequences of all 35 RAS subfamily GTPases and considered those with similar switch I regions to HRAS but that diverged in the switch II region (around HRAS Tyr64 and Met67 that contact RASSF5; **Figure S5**). Based on this, we selected six candidate GTPases: GEM, REM1, REM2, RASL12, ERAS and DIRAS3. These are highly conserved around RAS Asp33 and Asp38 but have distinct switch II sequences (**Figure 5B**). To assay interactions with RASSF1 we performed binding experiments using purified RA domain and activated GTPases. Mutational activation of full-length GTPases was based on RAS G12V, though the efficacy of this amino acid substitution in all small GTPases must by interpreted with caution (44). GST-tagged RASSF1 RA domain was incubated with cell lysates expressing the six activated GTPases or KRAS as a control. Remarkably, we found that all six GTPases could complex with the RASSF1 RA domain while KRAS did not (**Figure 5C**). This was corroborated by co-IP experiments in which FLAG-tagged RA domain (**Figure 5D**) or full-length RASSF1A (**Figure 5E**) were co-expressed with the GTPases. Precipitation with anti-FLAG revealed all six candidate GTPases complex with RASSF1. We consistently noted the RGK family proteins GEM, REM1 and REM2 as strong interactors, but could detect no binding to the fourth RGK GTPase, RRAD (not shown). Thus, a structure and sequence-based informatic approach was able to identify six potential RAS family GTPase partners for RASSF1.

**Figure 5.**
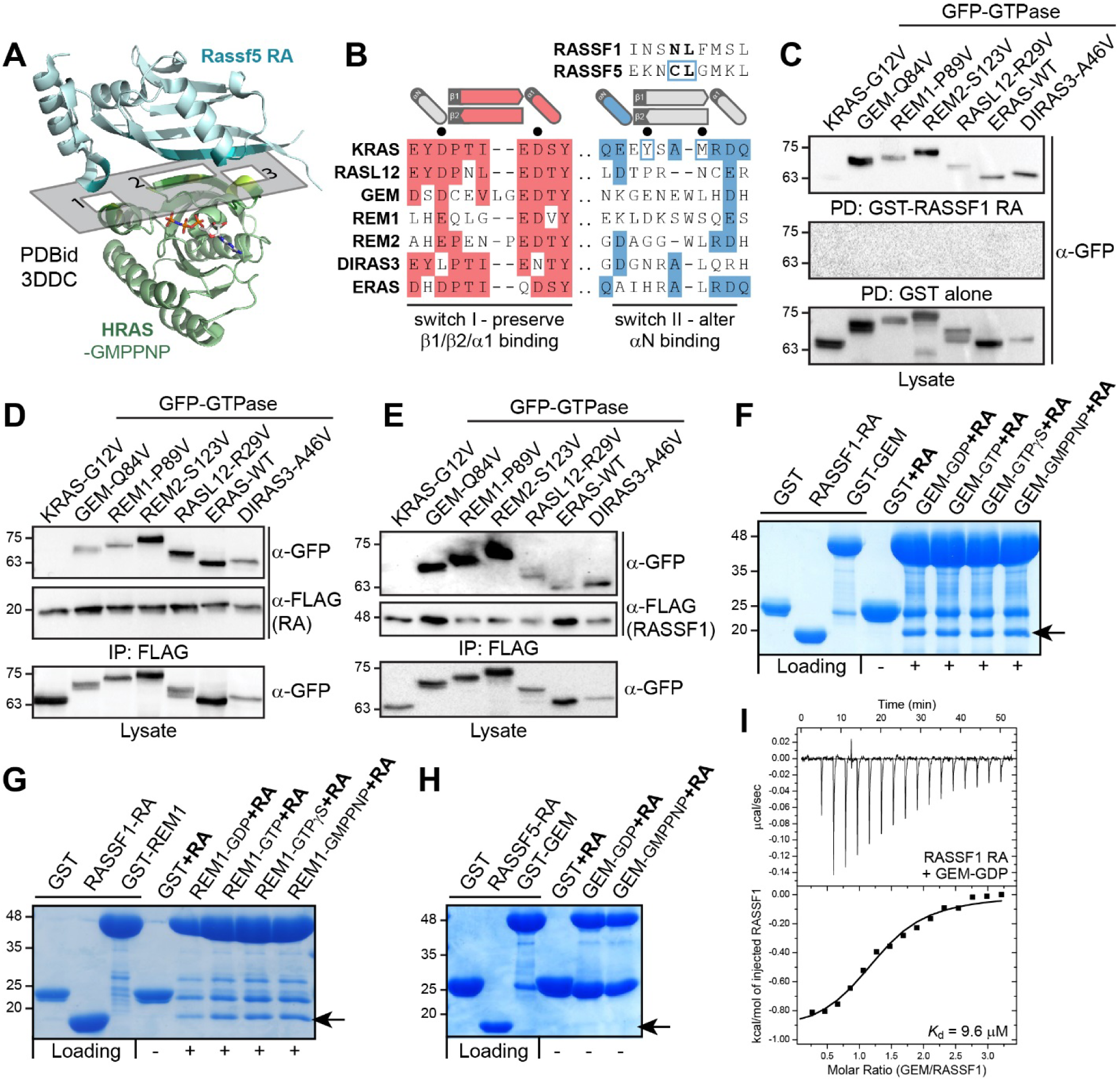
RASSF1A interacts with several alternative RAS subfamily GTPases. **(A)** Ribbons representation of the RASSF5-HRAS structure (PDBid 3DDC) reveals three non-linear interaction hot spots between RASSF5 and HRAS. **(B)** Amino acid sequence alignment of KRAS and the six candidate GTPase interactors GEM, REM1, REM2, RASL12, DIRAS3 and ERAS (see also Figure S5). These have high sequence similarity in the switch 1 region (sites 2 and 3 from (A)) but are highly divergent through switch II (site 1 from (A)). Selected candidates preserve key interacting residues for the RASSF β1/β2/α1 interface (schematic at top) but diverge around the αN-helix binding site. **(C)** Interactions between the candidate small GTPases and RASSF1 RA domain. Cells expressing GFP-tagged, activated GTPase mutants were incubated with purified GST-RASSF1 RA domain. Following precipitation on glutathione beads, GTPases interacting with RASSF1 were identified by anti-GFP immunoblot (top). The same lysates were incubated with GST alone as a control. **(D)** Complex between the candidate GTPases and RASSF1 determined by co-IP. Cells were co-transfected with vectors encoding GFP-tagged, activated GTPases and FLAG-RASSF1 RA domain. Anti-FLAG immunoprecipitation followed by anti-GFP immunoblot revealed interacting proteins (top). **(E)** Association of full-length RASSF1A with the candidate GTPases. Cells were co-transfected with vectors expressing GFP-GTPases and full-length FLAG-RASSF1A. Anti-GFP immunoblots of anti-FLAG immunoprecipitations reveal GTPases interacting with RASSF1A. **(F)** Direct binding of the GTPase GEM to the RASSF1 RA domain. Purified RA domain was incubated with purified, GST-tagged GEM exchanged with the nucleotides GDP, GTP, GTPγS or GMPPNP. Complexes were isolated on glutathione beads and washed 3 times. Arrow denotes the RA domain. No binding was observed to a GST alone control. **(G)** Direct binding of the RASSF1 RA domain to REM1. RA domain was incubated with purified, GST-REM1 loaded with GDP, GTP, GTPγS or GMPPNP. Complexes were isolated on glutathione beads and washed 3 times. Arrow denotes the RA domain. No binding was observed to GST alone. **(H)** No interaction was observed between the purified RASSF5 RA domain and either GDP- or GMPPNP-loaded GEM. **(I)** ITC analysis of the RASSF1 RA domain interaction with GEM. GDP-loaded GEM interacted with the RASSF1 RA domain with an affinity of 9.6 μM. The complex between RASSF1 and GTP-GEM proved highly unstable in solution and could not be measured.

To confirm these proteins directly interact we attempted *in vitro* mixing assays using purified RA domain with the RGK GTPases or RASL12. Unlike most RAS GTPases we’ve previously isolated, these four proteins were highly unstable under multiple expression and purification conditions. GEM and REM1 proved most stable in solution and were chosen for further study. GST-tagged, wild-type GEM (residues 62-203) and REM1 (residues 76-251) were purified and exchanged with either GDP, GTP, or the non-hydrolyzable analogs GTPγS and GMPPNP. The GTPases had a propensity to precipitate during exchange reactions, possibly due to their low affinity for nucleotide (45). GEM and REM1 were therefore exchanged using a simplified procedure: incubating for two hours in a 10-fold molar excess of nucleotide and 20 mM MgCl_2_ (46). The proteins were then mixed with purified RA domain and complexes isolated on glutathione beads. We observed a strong interaction between RASSF1 and both GEM (**Figure 5F**) and REM1 (**Figure 5G**) that was largely nucleotide independent. This is consistent with previous speculation that RGK GTPases may not behave as canonical switches (44). As a control, and to explore specificity, we mixed GDP- or GMPPNP-exchanged GEM with the RA domain of RASSF5 but observed no binding (**Figure 5H**). We next attempted to determine an affinity for GEM-RASSF1 using ITC. Purified GEM exchanged with GTP analogs and complexed with RASSF1 RA domain was too unstable for ITC and could not be analysed. However, we were able to measure a *K*_d_ of 9.6 μM for the interaction between GEM-GDP and RASSF1 (**Figure 5I**), an affinity comparable to other effector-GTPase partners. To summarize, we determined that RASSF1 is an *in vitro* binding partner of six RAS-like GTPases and can speculate the interactions depend on multiple binding hot spots, analogous to the RASSF5-HRAS complex. The promiscuity of effectors for GTPases is not well explored, but PI3K binds 15 distinct RAS family members (15) and we hypothesize most effectors will have several unique GTPase partners. Thus, any of the six RASSF1-interacting GTPases may be functional *in vivo* given the right context.

### RASSF1 and Candidate GTPase Binding Partners in Cells

Elucidating a functional role for the identified RASSF1-GTPase complexes is challenging. There are no known guanine nucleotide exchange factors (GEFs) for any of the six small GTPases, and there are no stimulatory conditions under which the endogenous GTPases are known to switch on. We began by resolving if mutationally activated GTPases could co-localize with RASSF1 in cells. Previous work has shown that RASSF1A localizes to microtubules through Microtubule Associated Proteins (MAPs) (47–49). Other data suggest RASSF1 isoforms are differentially localized and that RASSF1 localization is dynamic and dependent on the cell cycle (50–52). In HeLa cells we observed FLAG-tagged RASSF1A localized to microtubules. This localization was not dependent on its RA domain, which alone appeared cytoplasmic and enriched around the endoplasmic reticulum (ER) (**Figure 6A**). To determine if the GTPase partners co-localize with RASSF1, we had to consider their spatial restriction to the plasma membrane *via* lipidation. RASL12 and the RGK GTPases GEM, REM1 and REM2 completely lack predicted lipidation motifs, while DIRAS3 and ERAS are predicted to be farnesylated (**Figure 6B**). Expression of GFP-tagged, mutationally activated GTPases in HeLa cells verified that DIRAS3 and ERAS are plasma membrane-localized, like KRAS, while RASL12 was localized diffusely throughout the cell. Despite the lack of predicted lipidation, GEM, REM1 and REM2 were markedly associated with plasma membrane, as has been observed previously (53) (**Figures 6C** and **S6A**). When full-length RASSF1A was co-expressed with these GTPases we observed a striking recruitment of the only cytoplasmic GTPase, RASL12, to microtubules (**Figures 6D** and **S6A**). We did not observe significant fractions of the other GTPases on microtubules, consistent with their membrane-restriction. There was, however, a clear propensity for the RGK GTPases to cluster around regions of RASSF1A staining. Thus, RASSF1 can recruit RASL12 to microtubules, but its cytoskeletal attachment in HeLa cells precludes interaction with PM-associated GTPases.

**Figure 6.**
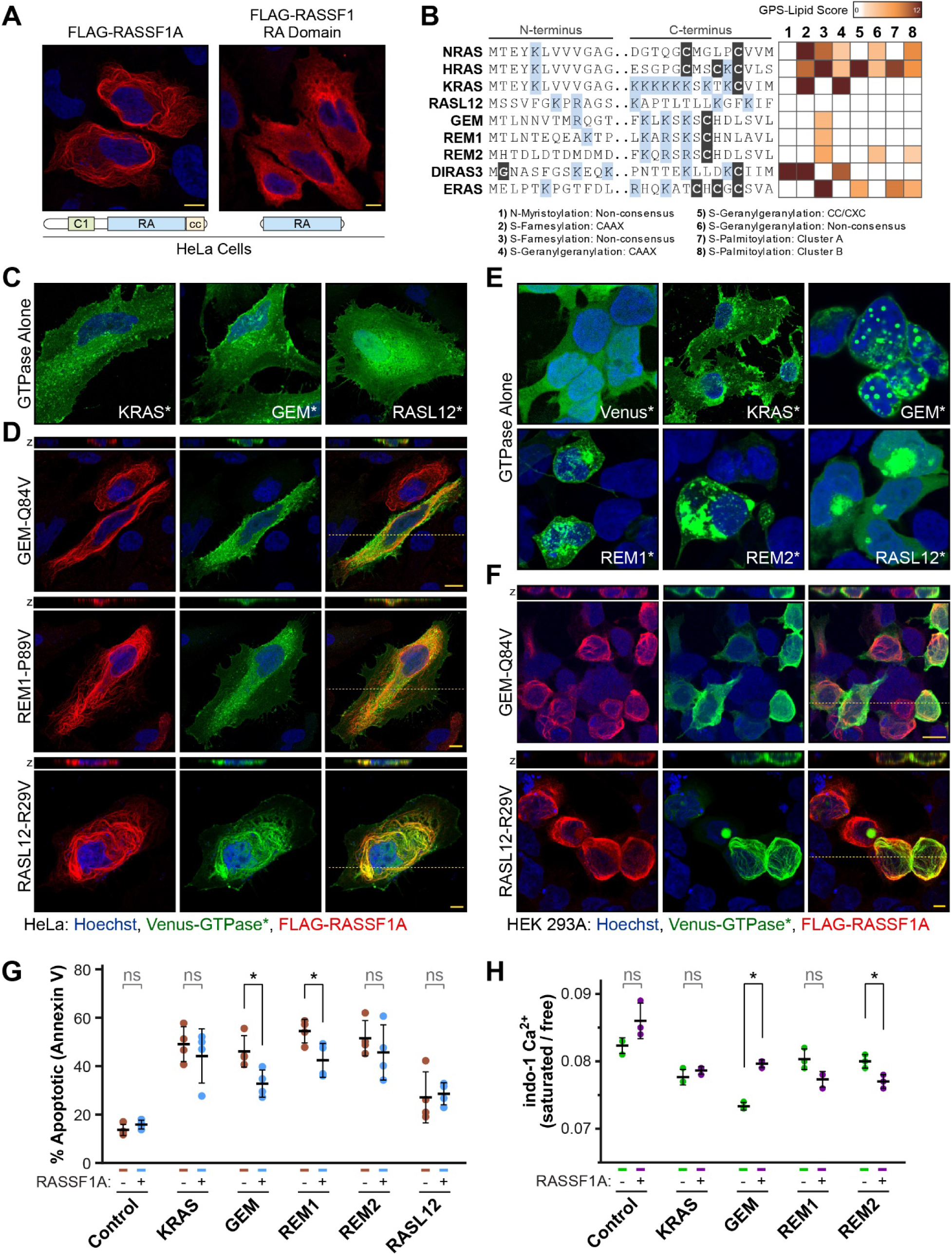
RASSF1 and interacting RAS family GTPases in cells. **(A)** Confocal microscopy shows the localization of full-length FLAG-RASSF1A (left) or the RASSF1 RA domain alone (right) in HeLa cells. **(B)** Prediction of lipidation sites in the candidate GTPases. Highlighted on an amino acid sequence alignment of the GTPase N- and C-termini regions are potential myristoylation or farnesylation sites (black boxes), as well as positively charged Lys and Arg residues (blue boxes). GPS-Lipid scores of potential lipidations are shown in the heat map, at right, colored from a low probability (white) to a high probability (dark brown). **(C)** Localization of GTPases expressed alone in HeLa cells, as determined by confocal microscopy. KRAS and the RGK GTPase GEM localize to the plasma membrane, while RASL12 has a diffuse localization throughout the cell. Asterisk’s indicate these are mutationally activated GTPases (based on RAS G12V). **(D)** Localization of the GEM, REM1 and RASL12 GTPases in HeLa cells when co-expressed with RASSF1A. Image shows the localization of mutationally activated, GFP-tagged GTPases (green) and FLAG-tagged RASSF1 (red). The *z*-plane is shown above each image, at a location indicated by the dashed line in the overlay. **(E)** Differential localization of GTPases in HEK 293A cells as detected by confocal microscopy. Asterisk’s indicate these are mutationally activated GTPases. **(F)** Co-expression of RASSF1A with these GTPases relieves puncta formation. Images show the localization of GFP-tagged GTPases (green) when co-expressed with FLAG-tagged RASSF1A (red) in HEK 293A cells. The *z*-plane is shown above each image at a location indicated by the dashed line. **(G)** Apoptotic induction by GTPases GEM, REM1, REM2 and RASL12. Annexin V staining and FACS analysis shows a large percentage of RGK expressing cells are undergoing apoptosis. Co-expression with RASSF1A significantly inhibits apoptosis in activated GEM- and REM1-expressing cells (asterisk indicates p<0.03). **(H)** Intracellular Ca^2+^ measured by Indo-AM staining and FACS analysis. Indo-AM loaded cells transfected with RGK GTPases have reduced intracellular Ca^2+^. Co-expression with RASSF1A significantly rescues this phenotype, while it is amplified when RASSF1A is co-expressed with REM2 (asterisk indicates p<0.025).

When we expressed activated RGK GTPases in HEK 293A cells they exhibited a completely altered pattern of localization. GEM formed large puncta throughout the cell, while REM1, REM2 and RASL12 aggregated in large, mostly cytoplasmic regions (**Figures 6E** and **S6B**). When the GTPases were co-expressed with full-length RASSF1A, these puncta completely disappear (**Figure 6F**). In these cells, RASSF1A and GTPases are co-distributed on microtubules and in the cytoplasm. These data suggest a functional link between RASSF1 and these RAS-like GTPases and highlight complexities to their direct association.

### RGK GTPases and RASSF1A Influence the Apoptotic Response and Ca^2+^ Homeostasis

There is not a single paper describing a function for RASL12, while numerous studies have attributed a role for RGK GTPases in regulating voltage-dependent Ca^2+^ channels. As most studies link RASSFs to apoptosis, we first determined if RASL12 or the RGK family members induce apoptosis and whether expression of RASSF1 might be synergistic. We used HEK 293T cells as they do not express detectable levels of endogenous RASSF1 (**Figure S6C**). Annexin V staining revealed that GEM (3.4-fold), REM1 (4-fold) and REM2 (3.8-fold) generate a significant apoptotic response over controls, levels similar to KRAS, while RASL12 induced a modest increase (2-fold) (**Figures 6G** and **S6D**). Co-expression with RASSF1 produced a significant decrease in the capacity for GEM (2.1-fold) and REM1 (2.7-fold) to induce apoptosis, while KRAS, REM2 and RASL12 levels were not significantly altered. These results suggest again that RASSF1 is not always a pro-apoptotic effector and may even have a protective function.

As RGK GTPases regulate intracellular Ca^2+^ levels we looked to establish a role for RASSF1 in Ca^2+^ signalling. We used the Indo-1 AM cell-permeant sensor to measure intracellular Ca^2+^ levels in cells expressing activated RGK GTPases alone, or with RASSF1A. Consistent with previous studies, we observed a significant reduction in free intracellular Ca^2+^ upon overexpression of GEM, and minor decreases when KRAS, REM1 or REM2 were overexpressed (**Figures 6H** and **S6E**). When RASSF1A was co-expressed with GEM, Ca^2+^ levels returned to near-normal levels. Conversely, co-expression of RASSF1A with REM1 or REM2 augmented their ability to restrict Ca^2+^ entry. RASSF1 expression had no impact on Ca^2+^ levels in cells expressing KRAS. Taken together, these data link RASSF1-RGK GTPase interactions with Ca^2+^ homeostasis, itself a master regulator of apoptosis and cell death, and begin to elucidate a functional role for the interaction of RASSF1 with the candidate GTPases.

### YAP1 nuclear localization is inhibited by RGK GTPases and RASSF effectors

Having established that RASSF1 complexes with both RGK GTPases and MST1 we next considered if the GTPases could regulate Hippo signalling. Immunofluorescent detection of endogenous YAP1 in U2OS cells shows its prominent nuclear localization when cells are sparsely distributed (Hippo off), and sequestration of YAP1 to the cytoplasm in packed cells (Hippo on; **Figure S7A**). We thus transfected GFP-tagged, activated RGK GTPases into U2OS cells plated at low density and monitored YAP1 localization. Expression of both GEM and REM1 resulted in abnormally elongated cells that markedly lacked nuclear YAP1, while KRAS had no effect (**Figure 7A**). There was a modest but significant reduction of nuclear YAP1 in cells expressing REM2, while YAP1 remained nuclear in the presence of RASL12 (**Figure S7B**). This suggests that RGK GTPases can activate Hippo signalling and sequester YAP1 in the cytoplasm.

**Figure 7.**
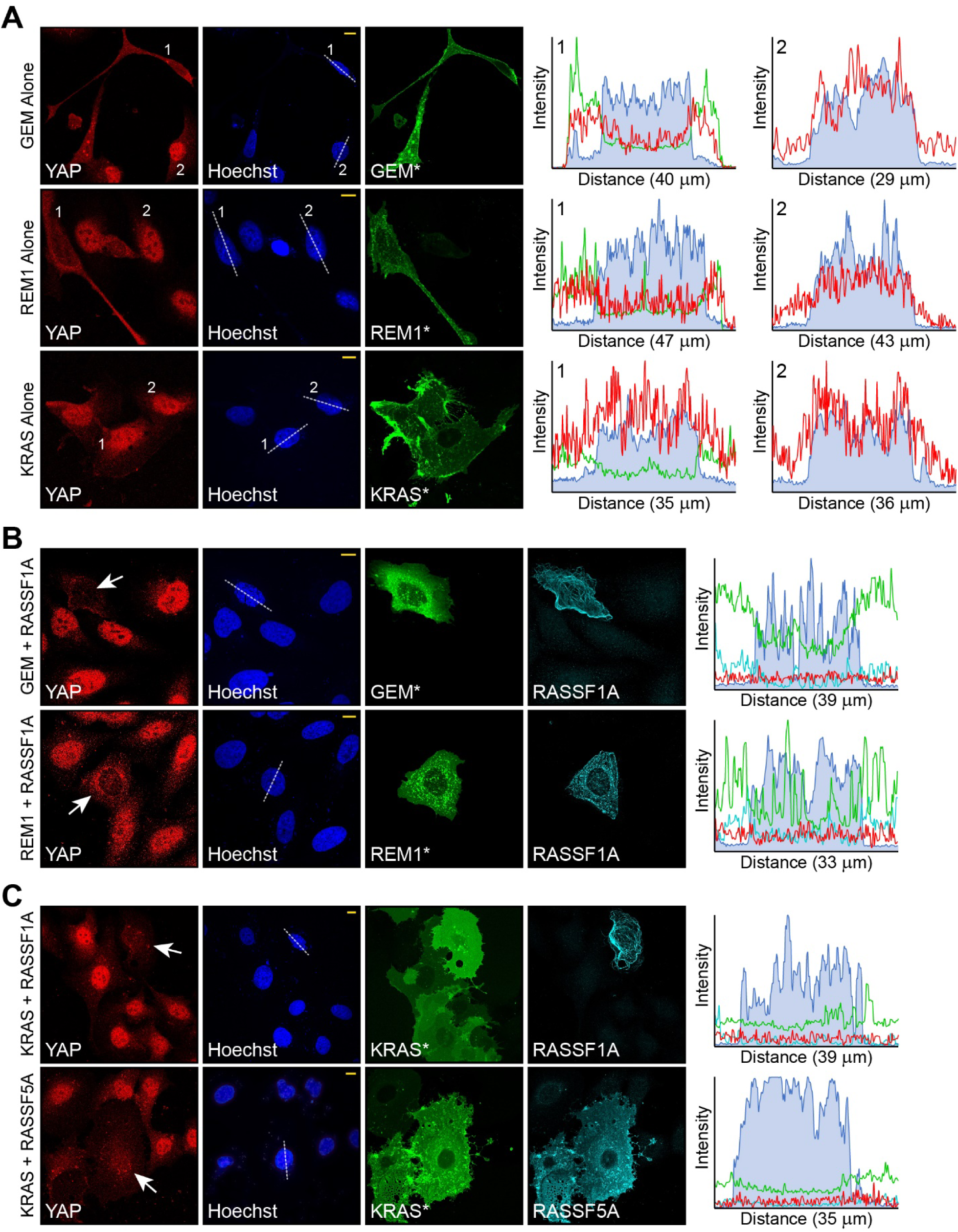
Hippo activation by RGK GTPases and RASSF effectors. **(A)** Confocal images show localization of endogenous YAP1 in U2OS cells expressing GFP-tagged, activated RGK GTPases (GEM or REM1), or KRAS alone. RGK GTPases induce significant cellular elongation and outgrowth. Quantification (right) was performed using *z*-stack fluorescence profiles from a plane through the nucleus (dotted line in Hoechst panel). Nuclei are blue, GTPase green and YAP1 red on the intensity graphs. YAP1 is excluded from the nucleus in cells expressing RGK GTPases (marked with a 1) compared to untransfected cells (marked with a 2). KRAS did not impact YAP1 localization. **(B)** YAP1 in U2OS cells co-expressing the RGK GTPase GEM or REM1 with their binding partner RASSF1A (FLAG-tagged). Quantification by fluorescence profiles is at right. Cells are no longer elongated, and YAP1 remains outside the nucleus. **(C)** Impact of FLAG-tagged RASSF1A or RASSF5A on YAP1 localization in U2OS cells expressing GFP-tagged, oncogenic KRAS. The presence of either RASSF protein restricts YAP1 nuclear localization, not observed when G12V-KRAS is expressed alone. In all panels, scale bar (yellow, Hoechst) represents 10 μm.

The capacity for RASSF proteins themselves to regulate Hippo signalling has not been well explored. We did not observe a significant change in YAP1 localization following expression of FLAG-tagged RASSF1A or RASSF5A alone (**Figure S7C**). However, the elongated cell shape phenotype induced by REM1 or GEM was completely rescued by co-expression with RASSF1A, and YAP1 remained cytoplasmic in these cells (**Figure 7B**). This suggests the irregular cell shape alone was not responsible for Hippo activation. Finally, we sought to answer whether RASSF proteins could regulate Hippo signalling downstream of activated KRAS. We co-expressed FLAG-RASSF1A or RASSF5A with GFP-tagged G12V-KRAS and stained for endogenous YAP1. While neither RASSFs or KRAS alone significantly impacts YAP1 localization, either RASSF1A or RASSF5A co-expressed with KRAS completely blocked YAP1 from the nucleus (**Figure 7C**). Thus, even RASSF effectors that do not directly complex with KRAS have a capacity to stimulate Hippo signalling and inactivate YAP in the presence of activated KRAS.

## Discussion

The distribution of signals to multiple downstream effector pathways is a central component of models describing small G-protein function. The specificity of effectors for GTPases is generally inferred based on homology, but we show that not all RA domain proteins are binding partners of H/K/N-RAS. We determined only one of the pro-apoptotic RASSF effectors can directly complex with these archetypal GTPases, underscoring the need for more stringent biochemical evidence to establish genuine effectors for each RAS family GTPase.

There has been significant interest in RASSF function from both a signalling and clinical standpoint in the past two decades. They represent one of the most highly conserved RAS effector families with orthologs in all metazoans and even single cell choanoflagellates. Tunicates have one N-RASSF ortholog and two C-RASSFs, one clustering with human RASSF2/4/6 by homology and the other with RASSF1/3/5 (**Figure S1**). The RASSF family thus expanded significantly in vertebrates, and we show this was accompanied by numerous amino acid substitutions to presumed GTPase binding residues. It is remarkable that relatively minor amino acid changes can significantly alter GTPase specificity, as we’ve demonstrated with RASSF5 Cys225 and RASSF1 Asn149. Following this, it is unsurprising that most RASSF family members are not true RAS effectors. Their diverse αN, β1, β2 and α1 sequences suggest RASSF effectors evolved to recognize distinct GTPases within the RAS superfamily. We also reveal that N-RASSFs are more highly related to ASPP effectors and directly complex with these p53-regulatory proteins. Despite this, there appears to be substantial functional overlap between C-RASSFs and N-RASSF/ASPPs, with evidence suggesting each of these scaffolds can sense changes in cell shape, adhesion or polarity to regulate senescence or apoptosis (35, 54–58). Indeed, all RASSF and ASPP family members were previously identified as network components that regulate cell growth in tandem with Hippo/STRIPAK, PP1 and polarity modules (59). It is intriguing to speculate that RASSFs may link the activation of diverse GTPases to these pathways.

There is a peculiar absence of small GTPases in proteomics networks. The explanation for this is likely complex, a combination of restricted expression, competition between effectors, and the requirement for high levels of the activated, GTP-bound conformation in assayed cells. We took a bioinformatic approach to resolve GTPase partners for the RASSF1 effector. As with other *RASSF* genes, expression of *RASSF1* is frequently silenced in numerous human cancers, and its tumour suppressor activity could be related to roles in autophagy (60), Hippo signalling (61–63), cell cycle regulation (50, 64), microtubule dynamics (47, 65), or apoptosis (48, 66). Though the dominant feature of the RASSF1 protein is its RA domain, its specificity for small G-proteins has not been investigated. Our results suggest that RASSF1 is an effector of the RAS subfamily GTPases GEM, REM1, REM2, RASL12, DIRAS3 and ERAS. We propose that most effectors bind multiple small GTPases rather than a single high affinity partner, and the context in which these specific GTPases are activated and bind RASSF1 must be determined. Also, our work does not exclude the possibility that RASSF1 could signal downstream of H/K/N-RAS, as it forms a heterodimer with RASSF5 *via* a SARAH domain interaction (67). Nevertheless, future studies should focus on the capacity for RASSF1 to signal from these understudied GTPases and elucidate how these interactions affect morphogenesis and cellular transformation.

The defined function of RGK GTPases is in regulating voltage-dependent Ca^2+^ channels and cell shape. There are multiple data linking RASSFs to Ca^2+^ homeostasis, including a functional interaction between RASSF1 and the Ca^2+^ pump PMCA4B (68) and a role for RASSF4 in regulating store-operated Ca^2+^ entry at ER–PM junctions (69). Indeed, fluctuations in intracellular Ca^2+^ upon RASSF overexpression may rationalize observations relating to apoptotic induction, as intracellular Ca^2+^ flux can dramatically influence apoptosis. To date, there is no available literature describing a functional role for the RASL12 GTPase, which we show is cytoplasmic and recruited to microtubules by RASSF1A. This protein clusters by homology with the RAS-like GTPases RERG, RASL11A/B and RASL10A/B (**Figure S5**) for which there are also no functional data (despite high evolutionary conservation). Implication of their interaction with RASSF1 thus remain to be elucidated, but the little data available suggest these small G-proteins also have a tumour suppressor role in human cancers (9, 70, 71).

While we did not observe an intrinsic ability for RASSFs to simulate apoptosis, either in the absence or presence of activated RAS, this does not preclude a role. Previous data linking RASSFs to apoptotic pathways suggest this association is complex. As with most Hippo network proteins, further study of RASSFs and apoptosis should employ cells or tissues in native, 3D environments. The strongest evidence for a RASSF-apoptosis connection derives from such studies (19), while those in 2D culture have generally relied on external stimuli to provoke a response (anti-FAS, Trail, etoposide, TNF, staurosporine). This also underscores the strong induction of apoptosis in 2D cultured cells induced by oncogenic RAS, documented here and by many others. We can now postulate this is not generated by a RAS-RASSF-Hippo signalling axis, but more likely *via* activation of p53 or alternative pathways (72–75).

Numerous recent works suggest KRAS and YAP1 converge to drive oncogenesis. In fact, active YAP1 can bypass KRAS addiction (76) and is required for neoplastic proliferation of numerous RAS-driven cancers (77–82). Despite this, there is no compelling studies linking RAS to Hippo activation through RASSFs. We detect a near complete loss of nuclear YAP1 in cells co-expressing G12V-KRAS and either RASSF5 or RASSF1. This could explain the frequent inactivation of *RASSF* genes in human cancers, and the molecular mechanisms directing RASSF control of YAP1 localization warrant further study. This is also true of the RGK GTPases, which we demonstrate here are novel Hippo pathway activators.

We have shown how a family of RA domain proteins expanded and diversified in vertebrates. Conserved RASSF interactions with Hippo kinases or ASPP proteins suggest specific RA domain-GTPase interactions may feed into these pathways, which has implications for both normal development and cellular transformation. We require further studies linking the more obscure RAS-like GTPases with their cognate effectors to help elucidate the full complexity of RAS-effector signalling.

## Materials and Methods

### Constructs and Antibodies

Expression constructs generated and used in this study:

| Construct | Backbone | Tag | Region (AA) | Source | Gene ID |
| --- | --- | --- | --- | --- | --- |
| GST alone | pGEX4-T2 | GST |  |  |  |
| GST-RASSF1-RA | pGEX4-T2 | GST | 123-296 | Human | 11186 |
| GST-RASSF2-RA | pGEX4-T2 | GST | 1-271 | Human | 9770 |
| GST-RASSF3-RA | pGEX4-T2 | GST | 41-194 | Human | 283349 |
| GST-Rassf4-RA | pGEX4-T2 | GST | 1-269 | Mouse | 213391 |
| GST-RASSF5-RA | pGEX4-T2 | GST | 199-367 | Human | 83595 |
| GST-Rassf6-RA | pGEX4-T2 | GST | 33-312 | Mouse | 73246 |
| GST-RASSF8-RA | pGEX4-T2 | GST | 1-87 | Human | 11228 |
| GST-RASSF9-RA | pGEX4-T2 | GST | 1-125 | Human | 9182 |
| GST-Rassf10-RA | pGEX4-T2 | GST | 1-151 | Mouse | 78748 |
| GST-RASSF1-RA-N149C | pGEX4-T2 | GST | 131-288 | Human | 11186 |
| GST-RASSF1-RA- | pGEX4-T2 | GST | 131-288 | Human | 11186 |
| GST-RASSF5-RA-C225N | pGEX4-T2 | GST | 199-367 | Human | 83595 |
| RASSF1-N149D/L150D | pGEX4-T2 | GST | 131-288 | Human | 11186 |
| RASSF1- N149K/L150K | pGEX4-T2 | GST | 131-288 | Human | 11186 |
| RASSF1- N149A/L150A | pGEX4-T2 | GST | 131-288 | Human | 11186 |
| GST-GEM | pGEX4-T2 | GST | 62-203 | Human | 2669 |
| GST-REM1 | pGEX4-T2 | GST | 76-251 | Human | 19700 |
| FLAG-RASSF1 | pCDNA5-FLAG-5' | FLAG | full-length | Human | 11186 |
| FLAG-RASSF2 | pDEST-pcDNA5-FLAG-5' | FLAG | full-length | Human | 9770 |
| FLAG-RASSF3 | pDEST-pcDNA5-FLAG-5' | FLAG | full-length | Human | 283349 |
| FLAG-Rassf4 | pDEST-pcDNA5-FLAG-5' | FLAG | full-length | Mouse | 213391 |
| FLAG-RASSF5 | pDEST-pcDNA5-FLAG-5' | FLAG | full-length | Human | 83595 |
| FLAG-Rassf6 | pDEST-pcDNA5-FLAG-5' | FLAG | full-length | Mouse | 73246 |
| FLAG-RASSF8 | pDEST-pcDNA5-FLAG-5' | FLAG | full-length | Human | 11228 |
| FLAG-RASSF9 | pDEST-pcDNA5-FLAG-5' | FLAG | full-length | Human | 9182 |
| FLAG-Rassf10 | pDEST-pcDNA5-FLAG-5' | FLAG | full-length | Mouse | 78748 |
| FLAG-RASSF1-RA | pCDNA5-FLAG-5' | FLAG | 131-288 | Human | 11186 |
| GFP alone | pVenus-C1 | GFP |  |  |  |
| GFP-MST-1 | pVenus-C1 | GFP | full-length | Human | 6789 |
| GFP-KRAS | pDEST-pcDNA3-Venus-5' | GFP | full-length | Human | 3845 |
| GFP-KRAS-G12V | pDEST-pcDNA3-Venus-5' | GFP | full-length | Human | 3845 |
| GFP-HRAS | pDEST-pcDNA3-Venus-5' | GFP | full-length | Human | 3265 |
| GFP-HRAS-G12V | pDEST-pcDNA3-Venus-5' | GFP | full-length | Human | 3265 |
| GFP-RHEB | pDEST-pcDNA3-Venus-5' | GFP | full-length | Human | 6009 |
| GFP-RHEB-Q64L | pDEST-pcDNA3-Venus-5' | GFP | full-length | Human | 6009 |
| GFP-RAP1A | pDEST-pcDNA3-Venus-5' | GFP | full-length | Human | 5906 |
| GFP-RAP1A-G12V | pDEST-pcDNA3-Venus-5' | GFP | full-length | Human | 5906 |
| GFP-RAC1 | pDEST-pcDNA3-Venus-5' | GFP | full-length | Human | 5879 |
| GFP-RAC1-G12V | pDEST-pcDNA3-Venus-5' | GFP | full-length | Human | 5879 |
| GFP-CDC42 | pDEST-pcDNA3-Venus-5' | GFP | full-length | Human | 998 |
| GFP-CDC42-G12V | pDEST-pcDNA3-Venus-5' | GFP | full-length | Human | 998 |
| GFP-RAN | pDEST-pcDNA3-Venus-5' | GFP | full-length | Human | 5901 |
| GFP-RAN-G19V | pDEST-pcDNA3-Venus-5' | GFP | full-length | Human | 5901 |
| GFP-GEM-Q84V | pDEST-pcDNA3-Venus-5' | GFP | full-length | Human | 2669 |
| GFP-REM1-P89V | pDEST-pcDNA3-Venus-5' | GFP | full-length | Human | 19700 |
| GFP-REM2-S123V | pDEST-pcDNA3-Venus-5' | GFP | full-length | Human | 161253 |
| GFP-RASL12-R29V | pDEST-pcDNA3-Venus-5' | GFP | full-length | Human | 51285 |
| GFP-ERAS-WT | pDEST-pcDNA3-Venus-5' | GFP | full-length | Human | 3266 |
| GFP-DIRAS3-A46V | pDEST-pcDNA3-Venus-5' | GFP | full-length | Human | 9077 |
| HIS-ASPP1 | pcDNA3.1/V5-HIS | HIS | full-length | Human | 23368 |
| HIS-ASPP2 | pcDNA3.1/V5-HIS | HIS | full-length | Human | 7159 |

We thank Dr. Matilda Katan (UCL, London) for providing cDNAs encoding murine Rassf10 and human RASSF5, Dr. Xin Lu (Ludwig Institute, Oxford) for ASPP1 and ASPP2 expression constructs, Dr. Anne-Claude Gingras (LTRI, Toronto) for human RASSF1, RASSF2, RASSF9 and murine Rassf4 cDNAs, Dr. Vuk Stambolic (PMCC, Toronto) for GFP-RHEB, and Dr. Jean-François Côté (IRCM, Montréal) for cDNAs encoding human GEM, REM1, REM2, RASL12, DIRAS3 and ERAS.

Antibodies used in this study:

| Antibody | Source |
| --- | --- |
| anti-FLAG-mouse | Sigma - F3165 |
| anti-FLAG-rabbit | Sigma - F7425 |
| anti-GFP | Abcam - ab290 |
| anti-HIS | Thermo - HIS.H8 - MA1-21315 |
| Goat anti-Mouse IgG-Tx-Red | Sigma - SAB3701076 |
| Annexin-V-Alexa Fluor 350 | Life Tech - V13246 |
| anti-RASSF1 | Abcam - ab23950 |
| anti-RASSF1 | ATLAS Antibodies - HPA040735 |
| anti-YAP1 (63.7) | Santa Cruz biotechnology - 10199 |
| Goat anti-Rabbit IgG-Alexa Fluor-647 | Invitrogen - A21244 |

### Protein Expression and Purification

Glutathione *S*-transferase (GST)- or 6xHistidine (HIS)-tagged proteins were expressed in *E. coli* (BL21-DE3-codon+) cells grown in LB media at 37°C. Cells were induced with 250 μM IPTG (isopropyl-β-D-thiogalactopyranoside) at OD 0.7. Following induction, cells were shifted to 16°C for growth overnight. Harvested cells were lysed in buffer (20□mM Tris-HCl (pH 7.5),

150□mM NaCl, 10% glycerol, 0.4% NP-40, protease inhibitors (Roche), and either 1□mM dithiothreitol or 10□mM β-mercaptoethanol) and sonicated. Lysate was clarified by centrifugation and incubated with Ni-NTA (Qiagen) or glutathione (Amersham) resins at 4°C for 1-2 hours. Bound protein was eluted by direct thrombin cleavage (GST) or with elution buffer containing 250□mM imidazole (Bioshop) followed by thrombin cleavage (HIS) or 30 mM reduced glutathione. Eluted proteins were purified by size exclusion chromatography using either an S75 or S200 column (GE).

### Nucleotide Exchange and Isothermal Titration Calorimetry

For nucleotide exchange, purified KRAS was incubated with a 10-fold molar excess (nucleotide:protein) of GMPPNP (Sigma) along with 10 mM EDTA and calf intestinal phosphatase at 37°C for 10 min. 20 mM MgCl_2_ was added, samples were incubated on ice for 10 mins and then dialyzed in the compatible buffer. RGK GTPases repeatedly precipitated using this methodology, so exchange of these GTPases was done with a 2-hour incubation in a 10-fold molar excess of nucleotide (GMPPNP, GTPγS or GTP (Sigma)) and 20 mM MgCl_2_. GTPase-RA domain interactions were measured using a MicroCal ITC200 (Malvern). Stock solutions were diluted into filtered and degassed 20 mM Tris-HCl (pH7.5), 150 mM NaCl and 1 mM DTT. Experiments were carried out at 25°C. Heats of dilution were determined from control experiments in which domains were titrated into buffer alone. Data were fit using Origin 7 (MicroCal).

### Cell Culture, Co-immunoprecipitation and Western Blotting

HEK 293T cells (ATCC CRL-3216) were seeded in six-well plates (at 3×10^6^) and transiently transfected with polyethylenimine (PEI) or Jet Prime. At least 1000 ng of DNA (unless otherwise stated) were transfected for each condition. 48 hours after transfection, cells were harvested and lysed in buffer (20 mM Tris-HCl (pH 7.5), 150 mM NaCl, 10% glycerol, 1% Triton X-100, 1 mM DTT and protease inhibitor P8340) for 10 minutes. Lysates were clarified by centrifugation and supernatant incubated with pre-washed Protein-G Sepharose and immunoprecipitating antibody. After 1 hour of incubation, beads were washed 3 times with lysis buffer. Beads containing bound proteins were reconstituted in SDS-loading buffer, separated using SDS-PAGE and transferred to nitrocellulose membrane for Western blot analysis. Membranes were blocked with TBST containing 5% skim milk. Following blocking, primary antibodies were detected using HRP conjugated anti-mouse Ig or anti-rabbit Ig antibodies (1:10000). Membranes were revealed by ECL reagent (Bio-Rad) and detection done by a Bio-Rad ChemiDoc imaging system equipped with ImageLab software.

### GST-RA Pull Downs

Lysates from HEK 293T cells expressing the protein of interest were incubated with GST-RA domain fusion protein or GST alone for 1 hour. Glutathione beads were used to capture GST fusion proteins or GST alone. Beads were washed 5 times with lysis buffer (20 mM Tris-HCl (pH 7.5), 150 mM NaCl, 5 mM MgCl_2_, 10% glycerol, 1% Triton X-100, 1 mM DTT and protease inhibitor P8340), eluted in SDS loading buffer and resolved by SDS-PAGE. Gels were transferred to nitrocellulose membrane and Western blot analysis performed as above.

### Microscopy

HeLa (ATCC CCL-2), HEK 293A (Thermo Fisher) or U2OS (ATCC HTB-96) cells were split in 6-well plates containing coverslips. 48 hours after transfection, cells were washed with phosphate buffered saline (PBS) and fixed with 4% paraformaldehyde (PFA). Fixed cells were permeabilized with 0.05% Tween or 0.1% Triton X100 (for U2OS cells) and blocked with 2% bovine serum albumin (BSA). Cells were incubated for 1 to 2 hours with primary antibody at 37°C or at RT (for U2OS cells) followed by secondary antibody and Hoechst staining. Finally, cells were treated with 70% then 95% ethanol and air dried before being mounted on glass slides. Slides were imaged using an LSM880 confocal microscope. Three lasers were used for detection (UV, Red and Far-Red). Images were processed using ZEN software.

### Structural Modelling

3D models for all RASSF (1-10) effectors were constructed using the automated protein structure homology model building program ‘MODBASE’ with energy minimization parameters available at the MODWEB server (https://modbase.compbio.ucsf.edu/modweb) (83, 84). Amino acid sequences based on our alignments were input in FASTA format. The server calculated structural models based on the best available template structures in the Protein Data Bank using MODPIPE, an automated modeling pipeline that relies on MODELLER for fold assignment, sequence-structure alignment, model building and model assessment (85). Models were evaluated for quality using Ramachandran plots and those with the highest DOPE score and fold reliability were selected for structural analysis.

### Measurements for Intracellular Ca^2+^ and Apoptosis

HEK 293T cells were seeded at a confluence of 0.4 million cells/ml/well in a 6 well plate. Following overnight incubation, cells were transfected with 1 μg of plasmid encoding GFP-tagged GTPases and/or FLAG-tagged RASSF along with controls using PEI. For measurements of intracellular Ca^2+^, 1 hour before harvesting cells they were incubated with 1 μM of Indo-1 AM at 37°C. Following this, cells were harvested by centrifugation and washed twice with PBS. Cells were analyzed using flowcytometry (YETI). In brief, cells were excited with a 350 nM laser and gated for an emission wavelength of 400 nM. Emission wavelength of the dye shifts from 475 nM in the Ca^2+^-free state to 400 nM when Ca^2+^-bound. Raw data from the Yeti was analyzed using Flow-Jo software. Apart form quantifying basal intracellular Ca^2+^ levels, Ca^2+^ levels after treatment with Ca^2+^ channel inhibitors (Thapsigargin) were also analyzed. For apoptosis: after 48 hours cells were harvested by trypsinization and dissolved in Annexin-V/Propidium Iodide (PI) buffer (10 mM HEPES (pH7.4), 150 mM NaCl, and 2.5 mM CaCl_2_ containing 2 μg of Annexin V 647-Fluor and PI). Samples were transferred to FACS tubes and analyzed using flowcytometry (YETI). Cells were sorted using sequential gating of Annexin, PI, and GFP channels. Raw data were analyzed using Flow-Jo software.

## Acknowledgements

This work was supported by grants (to M.J.S.) from the Canadian Institutes for Health Research (CIHR), the Canadian Cancer Society Research Institute (CCSRI) and the National Science and Engineering Council of Canada (NSERC). D.T. was supported by a scholarship from the Fonds de recherche du Québec-Nature et technologies (FRQNT). R.K. was supported by a research fellowship from the Fonds de recherche du Québec-Santé (FRQS). M.J.S. holds a Canada Research Chair in Cancer Signalling and Structural Biology.

## Author Contributions

D.T. and M.J.S. designed experiments. D.T., S.S., R.K., X.X. and M.J.S. performed experiments and analyzed data. A.S. and J.S. performed structural modelling. J.S. and M.J.S. supervised studies. D.T. and M.J.S. wrote the manuscript with input from R.K. and J.S.

