## Supplemental Data for "RASSF effectors couple diverse RAS subfamily GTPases to the Hippo pathway"

**Supplemental Information**

### Supplemental Figures

*Figure S1*

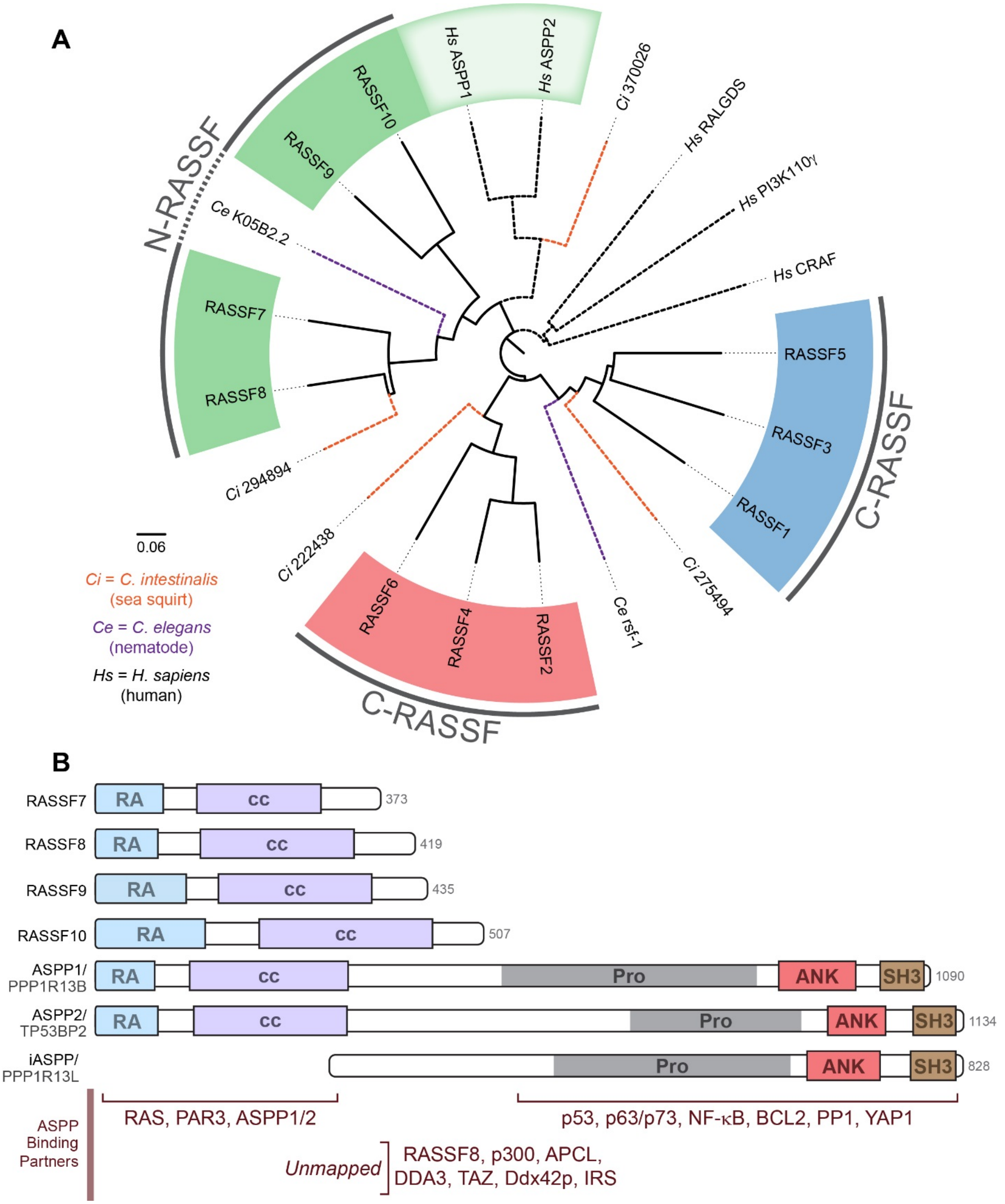

**Figure S1. Phylogenetic tree and domain architecture of RASSF proteins.**

**(A)** Phylogenetic relationship between human RASSF proteins, RASSF orthologs in *C. elegans* and *C. intestinalis*, and RA domains from several alternative human RAS effectors. The alignment is based on amino acid sequence conservation of the individual RA domains alone. C-RASSF1/3/5 cluster into one subgroup and C-RASSF2/4/6 into another. RA domains from N-RASSF7-10 are more highly related to those of the ASPP1/2 proteins. Nematodes have single N-RASSF and C-RASSF orthologs but no ASPP protein (APE-1 lacks an RA domain). Sea squirts, a close relative to the ancestral precursor of vertebrates, have single orthologs of the ASSP and N-RASSF effectors. They also have two C-RASSF orthologs, one related to the 2/4/6 cluster and the other to the 1/3/5 cluster. **(B)** A domain architecture schematic reveals similarity between ASPP and N-RASSF effectors. RA domains are in blue. Predicted or known coiled-coil regions are in purple. The ankyrin repeats (red) and SH3 domains (brown) of the ASPP proteins are missing in N-RASSFs. Several interaction partners have been mapped to distinct regions of the ASPP effectors (bottom).

*Figure S2*

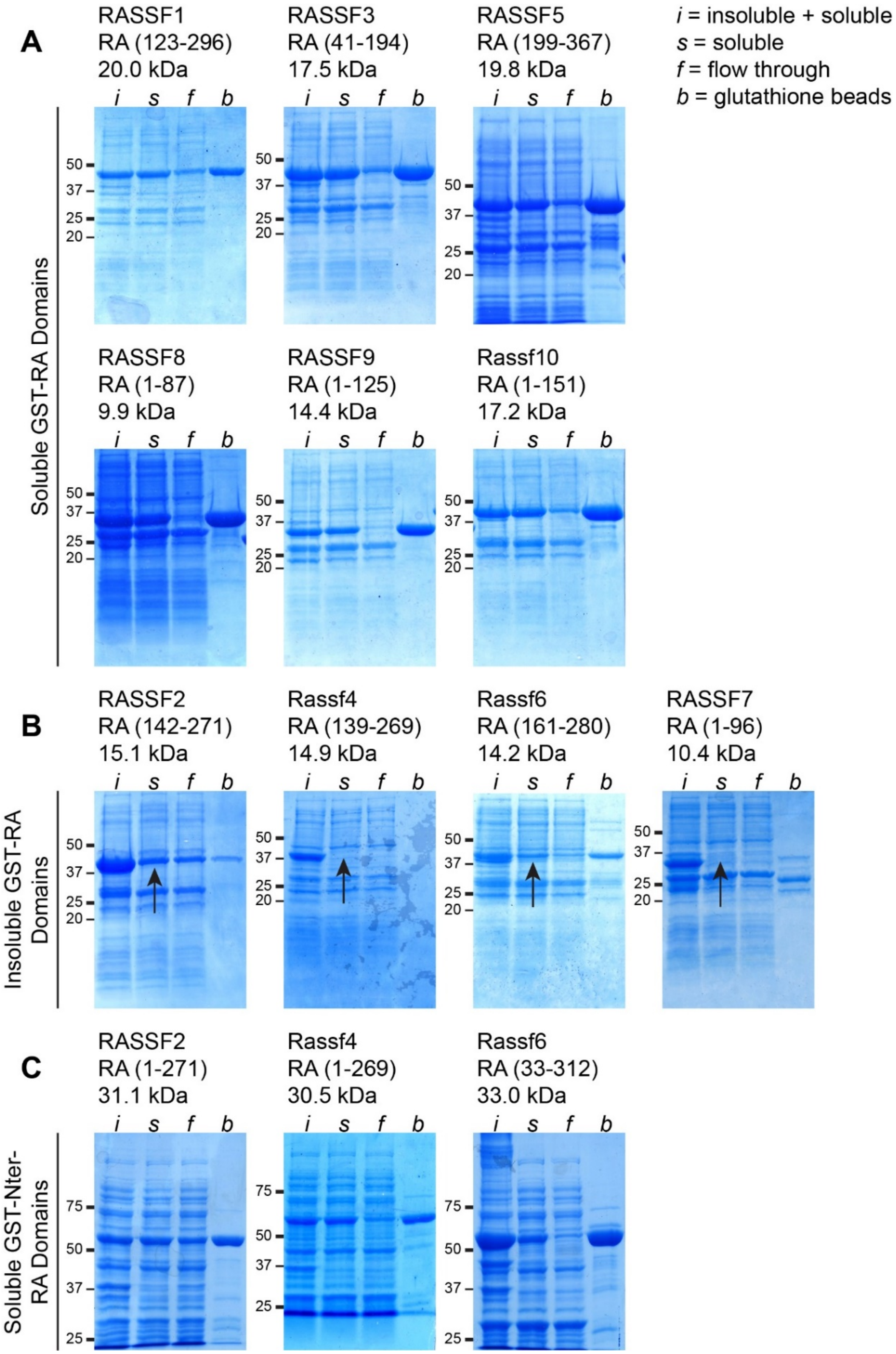
A
RASSF1
RA (123-296)
20.0 kDa
RASSF3
RA (41-194)
17.5 kDa
RASSF5
RA (199-367)
19.8 kDa
i = insoluble + soluble
s = soluble
f = flow through
b = glutathione beads
Soluble GST-RA Domains
RASSF8
RA (1-87)
9.9 kDa
RASSF9
RA (1-125)
14.4 kDa
Rassf10
RA (1-151)
17.2 kDa
B
RASSF2
RA (142-271)
15.1 kDa
Rassf4
RA (139-269)
14.9 kDa
Rassf6
RA (161-280)
14.2 kDa
RASSF7
RA (1-96)
10.4 kDa
Insoluble GST-RA Domains
C
RASSF2
RA (1-271)
31.1 kDa
Rassf4
RA (1-269)
30.5 kDa
Rassf6
RA (33-312)
33.0 kDa
Soluble GST-Nter-RA Domains
i s f b
75
50
37
25
20

**Figure S2. Purification of recombinant RASSF RA domains.**

**(A)** Six RASSF proteins could be expressed and purified to high concentration and high homogeneity using our initial domain boundaries (RASSF1, RASSF3, RASSF5, RASSF8, RASSF9 and RASSF10). Residues defining these boundaries and theoretical molecular weights of the resultant GST-fusion proteins are listed above each Coomassie stained gel. **(B)** Four RASSF proteins proved insoluble using our original boundaries (the related paralogs RASSF2, RASSF4 and RASSF6 as well as RASSF7). Domain boundaries and theoretical molecular weights are above each gel. Arrows indicate an absence of bands in the soluble fraction that were present in the insoluble/soluble fraction. **(C)** Incorporating the predicted secondary structure at the N-termini of RASSFs 2/4/6 significantly improved the solubility of these proteins. The amended domain boundaries and theoretical molecular weights are indicated above each Coomassie stained gel.

*Figure S3*

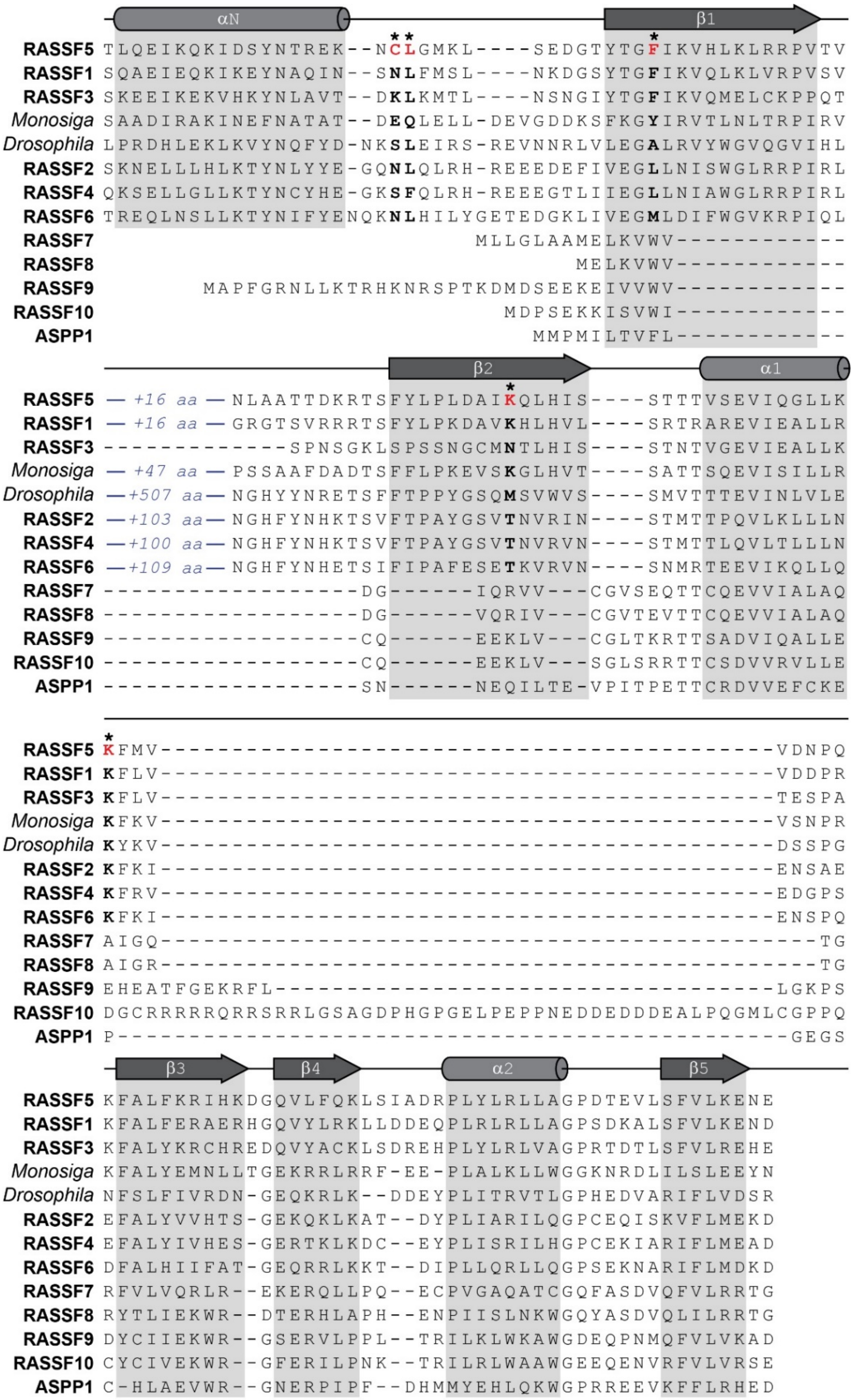

αN
β1
RASSF5 TLQEIKQKIDSYNTREK--NCLGMKL----SEDGTYTGFIKVHLKLRRPVTV
RASSF1 SQAEIEQKIKEYNAQIN--SNLFMSL----NKDGSYTGFIKVQLKLVRPVSV
RASSF3 SKEEIKEKVHKYNLAVT--DKLKMTL----NSNGIYTGFIKVQMELCKPPQT
Monosiga SAADIRAKINEFNATAT--DEQLELL-DEVGDDKSFKGYIRVTLNLTRPIRV
Drosophila LPRDHLEKLKVYNQFYD-NKSLEIRS-REVNNRLVLEGALRVYWGVQGVIHL
RASSF2 SKNELLLHLKTYNLYYE-GQNLQLRH-REEEDEFIVEGLLNISWGLRRPIRL
RASSF4 QKSELLGLLKTYNCYHE-GKSFQLRH-REEEGTLIIEGLLNIAWGLRRPIRL
RASSF6 TREQLNSLLKTYNIFYENQKNLHILYGETEDGKLIVEGMLDIFWGVKRPIQL
RASSF7 MLLGLAAMELKVWV-------------
RASSF8 MELKVWV-------------
RASSF9 MAPFGRNLLKTRHKNRSPTKDMDSEEKEIVVWV-------------
RASSF10 MDPSEKKISVWI-------------
ASPP1 MMPMILTVFL-------------
β2
α1
RASSF5 — +16 aa — NLAATTDKRTSFYLPLDAIKQLHIS----STTTVSEVIQGLLK
RASSF1 — +16 aa — GRGTSVRRRTSFYLPKDAVKHLHVL----SRTRAREVIEALLR
RASSF3 -------------SPNSGKLSPSSNGCMNTLHIS----STNTVGEVIEALLK
Monosiga — +47 aa — PSSAAFDADTSFFLPKEVSKGLHVT----SATTSQEVISILLR
Drosophila —+507 aa— NGHYYNRETSFFTPPYGSQMSVWVS----SMVTTTEVINLVLE
RASSF2 —+103 aa— NGHFYNHKTSVFTPAYGSVTNVRIN----STMTTPQVLKLLLN
RASSF4 —+100 aa— NGHFYNHKTSVFTPAYGSVTNVRVN----STMTTLQVLTLLLN
RASSF6 —+109 aa— NGHFYNHETSIFIPAFESETKVRVN----SNMRTEEVIKQLLQ
RASSF7 ------------------DG------IQRVV---CGVSEQTTCQEVVIALAQ
RASSF8 ------------------DG------VQRIV---CGVTEVTTCQEVVIALAQ
RASSF9 ------------------CQ------EEKLV---CGLTKRTTSADVIQALLE
RASSF10 ------------------CQ------EEKLV---SGLSRRTTCSDVVRVLLE
ASPP1 ------------------SN------NEQILTE-VPITPETTCRDVVEFCKE
RASSF5 KFMV-------------------------------------------VDNPQ
RASSF1 KFLV-------------------------------------------VDDPR
RASSF3 KFLV-------------------------------------------TESPA
Monosiga KFKV-------------------------------------------VSNPR
Drosophila KYKV-------------------------------------------DSSPG
RASSF2 KFKI-------------------------------------------ENSAE
RASSF4 KFRV-------------------------------------------EDGPS
RASSF6 KFKI-------------------------------------------ENSPQ
RASSF7 AIGQ----------------------------------------------TG
RASSF8 AIGR----------------------------------------------TG
RASSF9 EHEATFGEKRFL-----------------------------------LGKPS
RASSF10 DGCRRRRRQRRSRRLGSAGDPHGPGELPEPPNEDDEDDDEALPQGMLCGPPQ
ASPP1 P-----------------------------------------------GEGS
β3
β4
α2
β5
RASSF5 KFALFKRIHKDGQVLFQKLSIADRPLYLRLLAGPDTEVLSFVLKENE
RASSF1 KFALFERAERHGQVYLRKLLDDEQPLRLRLLAGPSDKALSFVLKEND
RASSF3 KFALYKRCHREDQVYACKLSDREHPLYLRLVAGPRTDTLSFVLREHE
Monosiga KFALYEMNLLTGEKRRLRRF-EE-PLALKLLWGGKNRDLILSLEEYN
Drosophila NFSLFIVRDN-GEQKRLK--DDEYPLITRVTLGPHEDVARIFLVDSR
RASSF2 EFALYVVHTS-GEKQKLKAT--DYPLIARILQGPCEQISKVFLMEKD
RASSF4 EFALYIVHES-GERTKLKDC--EYPLISRILHGPCEKIARIFLMEAD
RASSF6 DFALHIIFAT-GEQRRLKKT--DIPLLQRLLQGPSEKNARIFLMDKD
RASSF7 RFVLVQRLR--EKERQLLPQ--ECPVGAQATCGQFASDVQFVLRRTG
RASSF8 RYTLIEKWR--DTERHLAPH--ENPIISLNKWGQYASDVQLILRRTG
RASSF9 DYCIIEKWR--GSERVLPPL--TRILKLWKAWGDEQPNMQFVLVKAD
RASSF10 CYCIVEKWR--GFERILPNK--TRILRLWAAWGEEQENVRFVLVRSE
ASPP1 C-HLAEVWR--GNERPIPF--DHMMYEHLQKWGPRREEVKFFLRHED

**Figure S3. Sequence alignment of RASSF proteins.**

Multiple sequence alignment of all ten human RASSF RA domains along with human ASPP1 and RASSF orthologs from the fruit fly *D. melanogaster* and single cell choanoflagellate *M. brevicollis*. Secondary structure regions as derived from the crystal structure of RASSF5 bound to HRAS (PDBid 3DDC) are indicated at top. Amino acid conservation and secondary structure predictions reveal a high degree of homology in all domains from the α1-helix to the C-termini. The β1 and β2 strands of N-RASSFs 7-10 are shorter and they do not encode an αN-helix, analogous to human ASPP1. The RA domain of RASSF5 includes a large, unstructured loop of 29 residues between β1 and β2 that provided no electron density in the published crystal structure. This loop is a similar length in RASSF1 and shorter in RASSF3. The regions of RASSF2, RASSF4 and RASSF6 that are most homologous to the β1 and αN regions of RASSF5 are separated by long insertions of approximately 100 residues from their homologous β2 strands. The IUPred tool (https://iupred2a.elte.hu/) predicts these are intrinsically disordered regions and they are rich in Ser and Arg. This loop is over 50 residues in the lone *M. brevicollis* RASSF ortholog and has expanded to over 500 residues in the single *D. melanogaster* ortholog. Impacts on RA domain folding are unknown, but the domains are insoluble in the absence of the N-terminal homology regions. RASSF5 residues undergoing side chain interactions with HRAS (PDBid 3DD) are in bold red, marked with an asterisk. Corresponding residues in the aligned domains are in bold black.

*Figure S4*

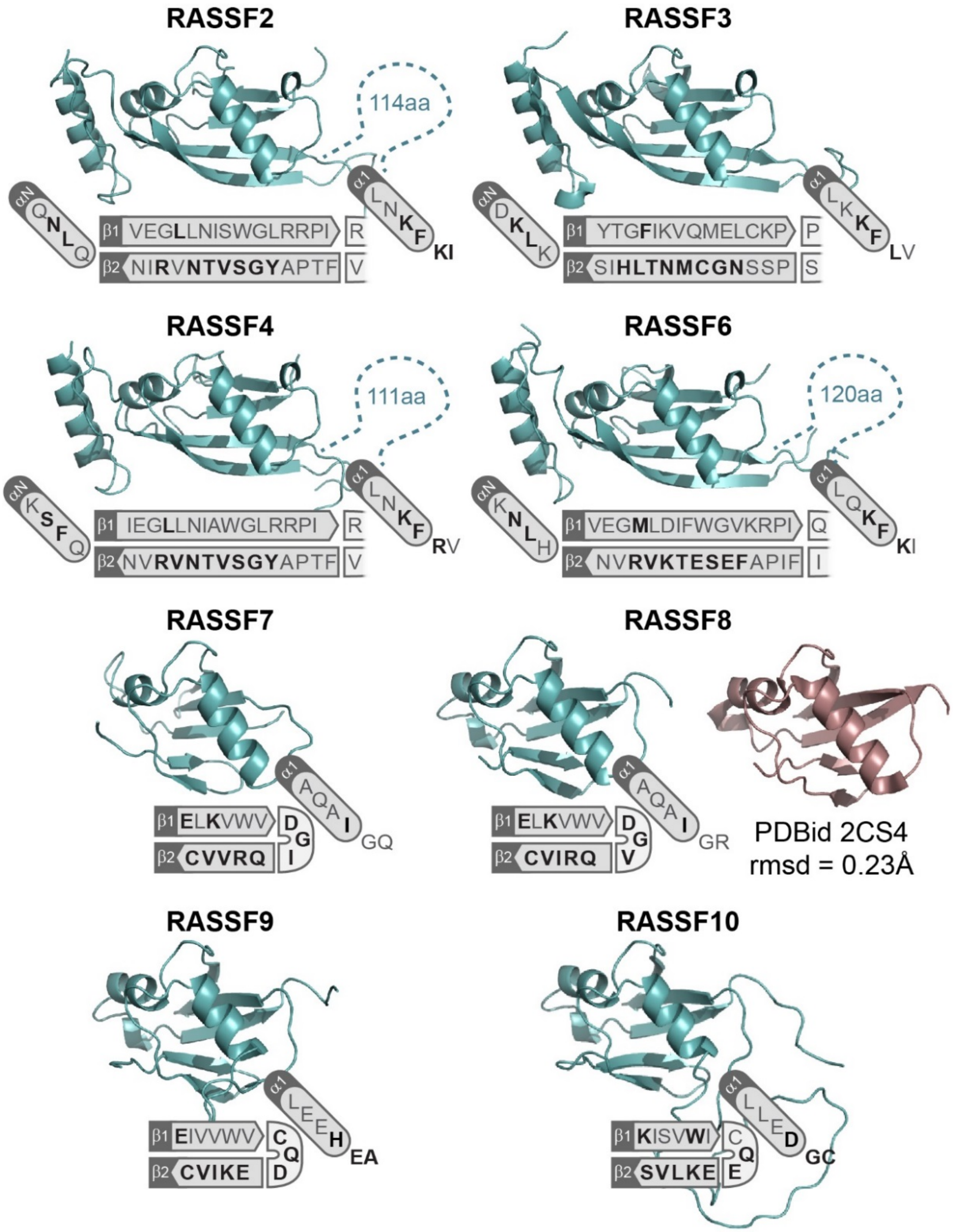
RASSF2
114aa
αN QNLQ
β1 VEGLLNISWGLRRPI R
β2 NIRVNTVSGYAPTF V
α1 LNKF KI
RASSF3
αN DKLK
β1 YTGFIKVQMELCKP P
β2 SIHLTNMCGNSSP S
α1 LKKF LV
RASSF4
111aa
αN KSFQ
β1 IEGLLNIAWGLRRPI R
β2 NVRVNTVSGYAPTF V
α1 LNKF RV
RASSF6
120aa
αN KNLH
β1 VEGMLDIFWGVKRPI Q
β2 NVRVKTESEFAPIF I
α1 LQKF KI
RASSF7
β1 ELKVWV D
β2 CVVRQ
G I
α1 AQAI GQ
RASSF8
β1 ELKVWV D
β2 CVIRQ
G V
α1 AQAI GR
PDBid 2CS4
rmsd = 0.23Å
RASSF9
β1 EIVVWV C
β2 CVIKE
Q D
α1 LEEH EA
RASSF10
β1 KISVWI C
β2 SVLKE
Q E
α1 LLED GC

**Figure S4. Structural models of RASSF RA domains.**

Ribbons representation of homology modelled RASSF RA proteins. Models of domains from C-RASSFs 2/3/4/6 were based on the RASSF5-HRAS structure (PDBid 3DDC). An AFDN-HRAS structure (PDBid 6AMB) served as a template for domains from N-RASSFs 7-10. A solution structure of the RASSF8 RA domain solved by NMR (PDBid 2CS4) has a backbone r.m.s.d. of just 0.23 Å compared with the modelled structure, validating the approach. A schematic showing amino acid positions in the β1 and β2 strands, and the αN (if present) and α1 helices is shown below each structure. Residues within 4 Å of the GTPase are in bold.

*Figure S5*

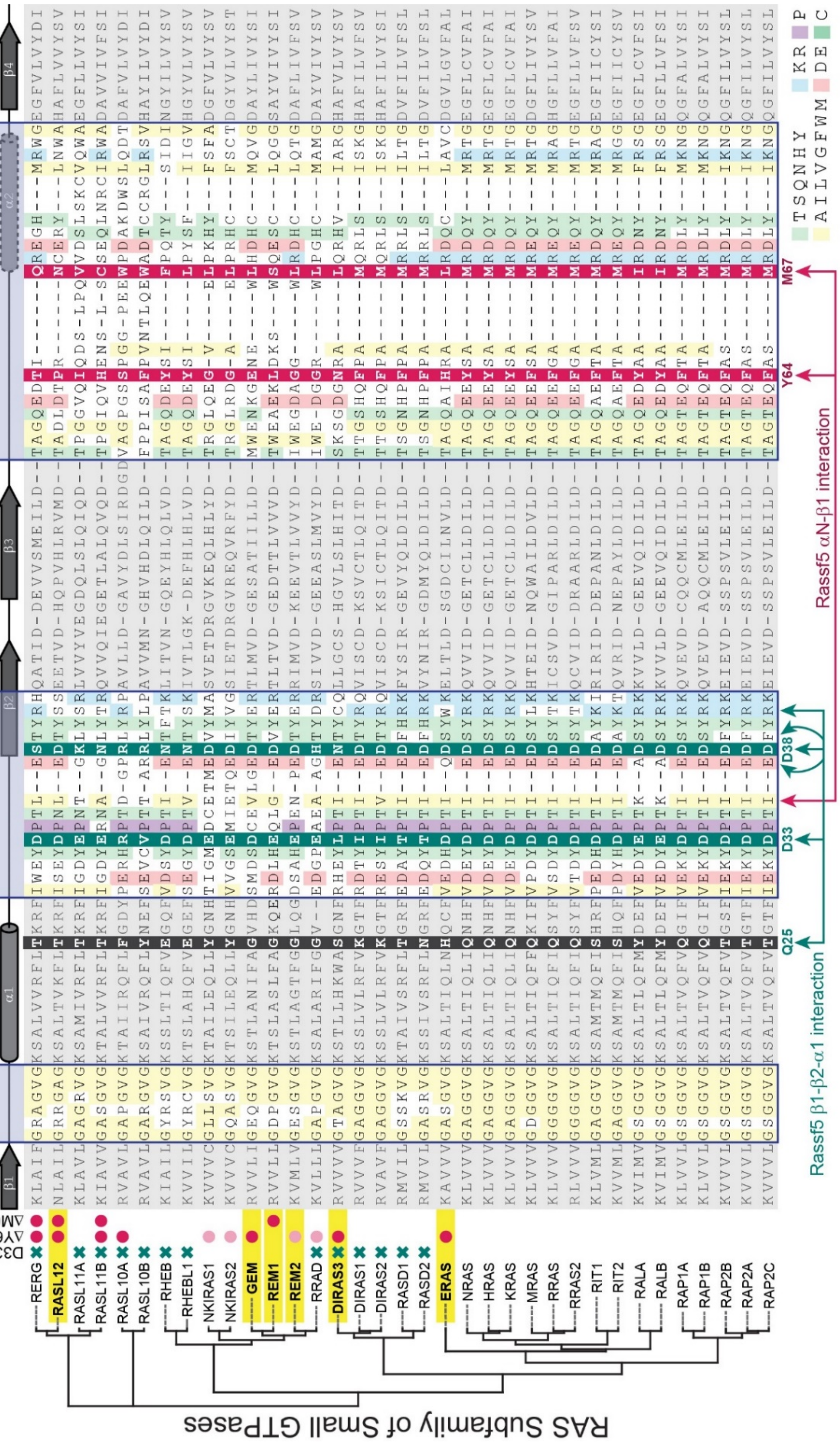
RAS Subfamily of Small GTPases
p-loop
switch I / effector
switch II
β1
α1
β2
β3
α2
β4
D33/38
ΔY64
ΔM67
Q25
D33
D38
Y64
M67
Rassf5 β1-β2-α1 interaction
Rassf5 αN-β1 interaction
RERG
RASL12
RASL11A
RASL11B
RASL10A
RASL10B
RHEB
RHEBL1
NKIRAS1
NKIRAS2
GEM
REM1
REM2
RRAD
DIRAS3
DIRAS1
DIRAS2
RASD1
RASD2
ERAS
NRAS
HRAS
KRAS
MRAS
RRAS
RRAS2
RIT1
RIT2
RALA
RALB
RAP1A
RAP1B
RAP2B
RAP2A
RAP2C
TSQNHY
AILVGFWM
KR
DE
P
C

**Figure S5. Identification of candidate GTPases in the RAS subfamily that could interact with RASSF1.**

Multiple sequence alignment of all 35 RAS subfamily small GTPases. Amino acid sequences starting just upstream of the p-loop and ending after the switch II region are shown. Regions of secondary structure are depicted at top. GTPases are ordered based on their phylogenetic relationship (tree at left). The p-loop, switch I/effector and switch II regions are highlighted while regions outside are in grey. GTPases with significant amino acid substitutions at the RAS Asp33 and Asp38 locations are marked with a green 'X' (left). Those with divergent switch II sequences around RAS Tyr64 or Met67 are marked with a red circle. As candidate RASSF1 interacting proteins we chose six GTPases that were conserved around Asp33 and Asp38 but diverged at Tyr64 and/or Met67: GEM, REM1, REM2, RALS12, ERAS and DIRAS3. These are highlighted yellow.

*Figure S6*

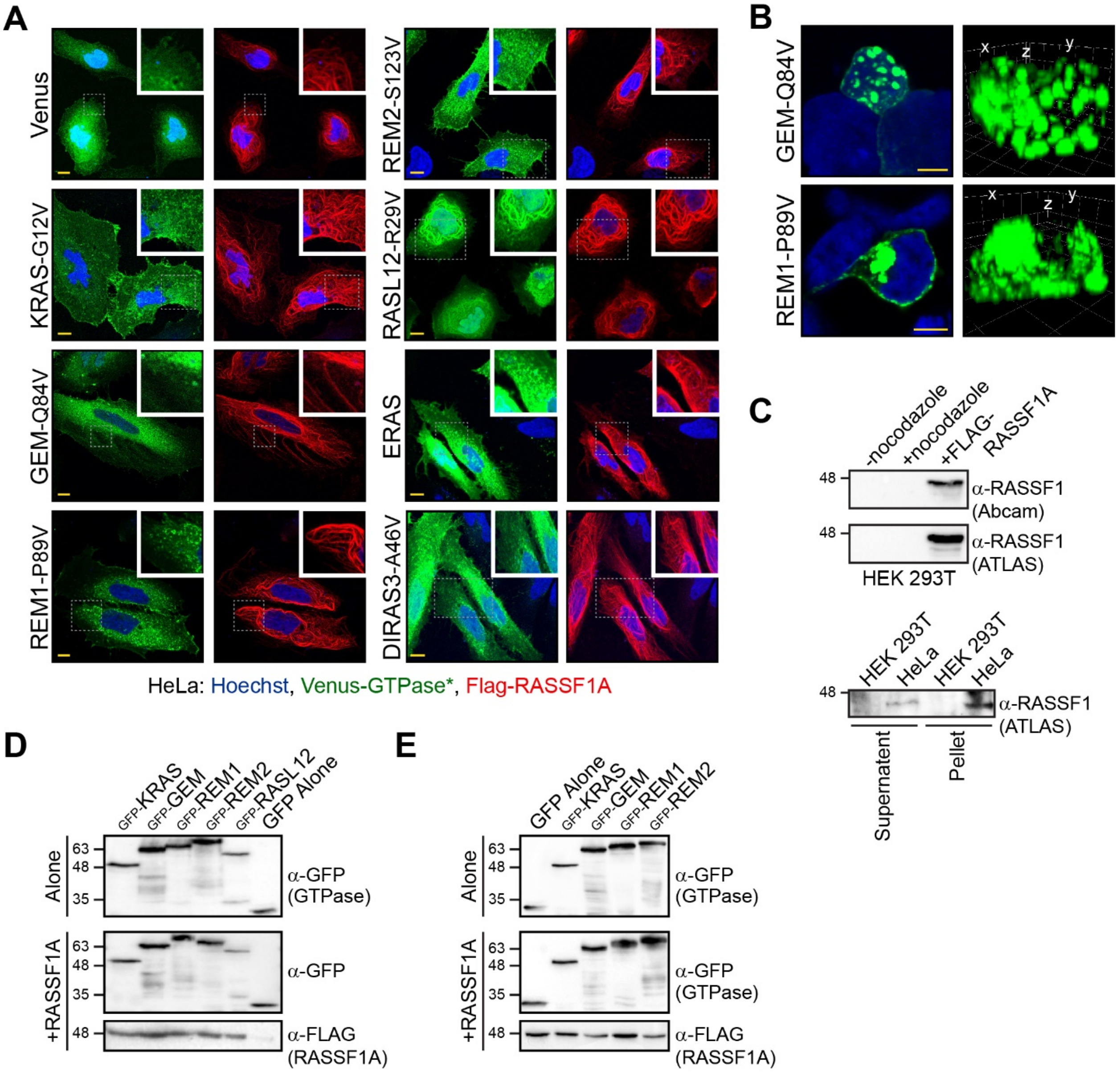
A
Venus
KRAS-G12V
GEM-Q84V
REM1-P89V
REM2-S123V
RASL12-R29V
ERAS
DIRAS3-A46V
HeLa: Hoechst, Venus-GTPase*, Flag-RASSF1A
B
GEM-Q84V
REM1-P89V
x
z
y
C
-nocodazole
+nocodazole
+FLAG-RASSF1A
48
α-RASSF1 (Abcam)
α-RASSF1 (ATLAS)
HEK 293T
HEK 293T
HeLa
α-RASSF1 (ATLAS)
Supernatent
Pellet
D
GFP-KRAS
GFP-GEM
GFP-REM1
GFP-REM2
GFP-RASL12
GFP Alone
Alone
+RASSF1A
63
48
35
α-GFP (GTPase)
α-GFP
α-FLAG (RASSF1A)
E
GFP Alone
GFP-KRAS
GFP-GEM
GFP-REM1
GFP-REM2
α-GFP (GTPase)
α-FLAG (RASSF1A)

**Figure S6. RASSF1 and candidate GTPases in cells.**

(**A**) Co-localization of GTPases and RASSF1 in HeLa cells as detected by confocal microscopy. Images show subcellular localization of GFP-tagged, activated GTPases (green) in the presence of FLAG-tagged RASSF1 RA domain (red, left) or full length RASSF1A (red, right). RASSF1 proteins were detected using anti-FLAG and a TxRed-conjugated secondary antibody. Insets are enlargements of the areas boxed in dotted lines. Scale bar (yellow) represents 10 µm. **(B)** Puncta distribution of activated GEM and REM1 in HEK 293A cells. Shown are both 2D (left) and 3D (right) renders of confocal images taken of GFP-tagged GEM and REM1. The *x*, *y* and *z* planes are labelled on the 3D render. These puncta are observed throughout the cell, while also in the nucleus in GEM-overexpressing cells. **(C)** HEK 293T cells do not express detectable levels of endogenous RASSF1. Two commercial antibodies against RASSF1 (purchased from Abcam and ATLAS Antibodies) were used to probe RASSF1 expression levels. No RASSF1 was present in HEK 293T cell lysates (+/- nocodazole to release protein from microtubules), but overexpressed RASSF1 was positively detected (top). In comparison, a small amount of endogenous RASSF1 was detected in HeLa cell lysates after long exposure (bottom), though much of this protein was found in the insoluble pellet. **(D)** A fraction of cells assayed for apoptotic activity were used to verify expression levels of FLAG-tagged RASSF and GFP-tagged GTPase proteins. Immunoblots with anti-FLAG and anti-GFP are shown for RGK assay**.** **(E)** A fraction of cells assayed for intracellular $Ca^{2+}$ entry were used to verify expression levels of FLAG-tagged RASSF1A and GFP-tagged GTPases. Immunoblots with anti-FLAG and anti-GFP are shown.

*Figure S7*

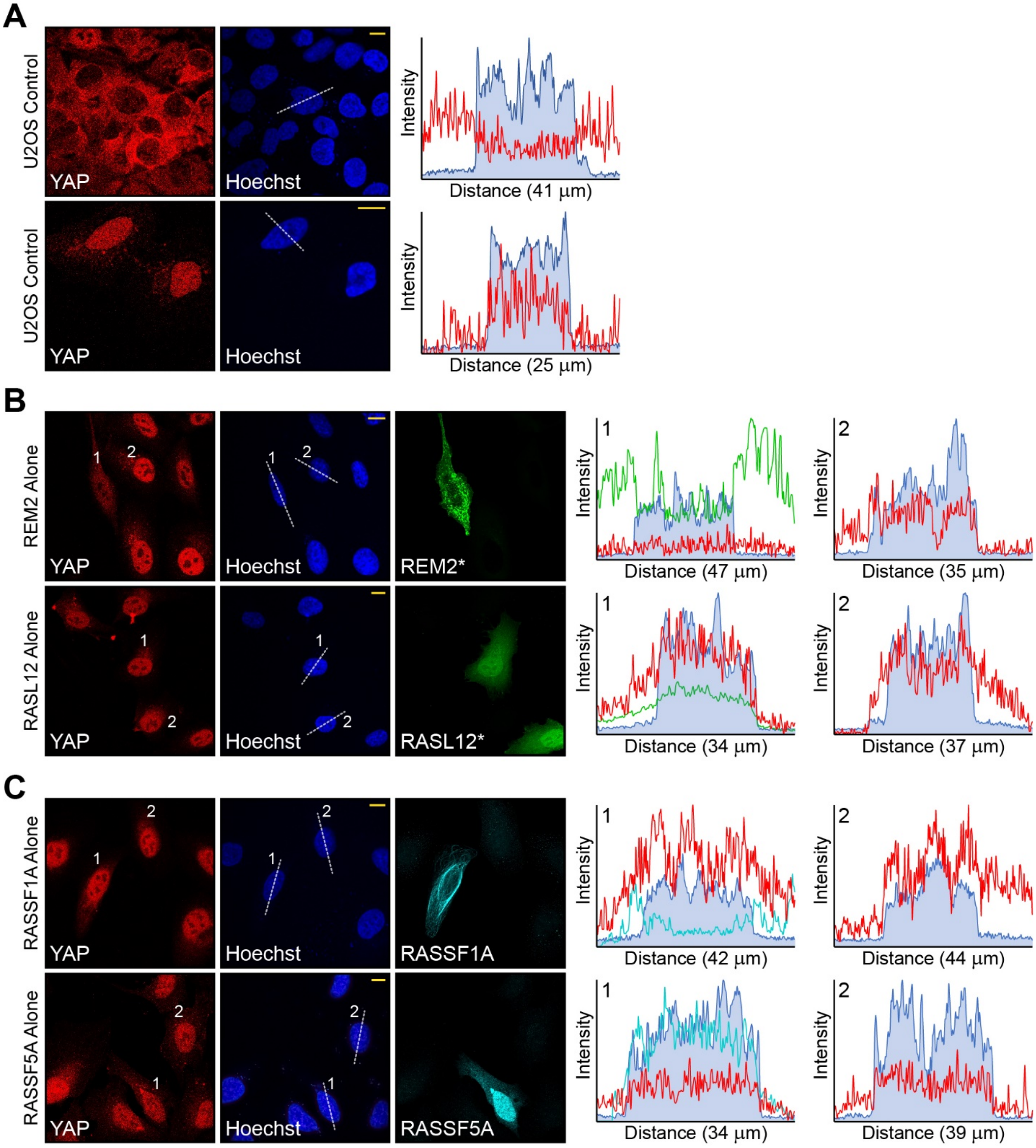
A
U2OS Control
YAP
Hoechst
Intensity
Distance (41 µm)
U2OS Control
YAP
Hoechst
Intensity
Distance (25 µm)
B
REM2 Alone
YAP
Hoechst
REM2*
Intensity
Distance (47 µm)
Distance (35 µm)
RASL12 Alone
YAP
Hoechst
RASL12*
Distance (34 µm)
Distance (37 µm)
C
RASSF1A Alone
YAP
Hoechst
RASSF1A
Distance (42 µm)
Distance (44 µm)
RASSF5A Alone
YAP
Hoechst
RASSF5A
Distance (34 µm)
Distance (39 µm)

**Figure S7. Localization of YAP1 in U2OS cells determines Hippo pathway activation.**

**(A)** Confocal microscopy shows the localization of endogenous YAP1 in U2OS cells (left). Quantification was performed using fluorescence profiles taken from a plane of *z*-stacks through the nucleus (marked with dotted line on Hoechst panel). On the intensity profiles, blue is from the Hoechst-stained nucleus and red from YAP1. In these cells, YAP1 is localized in the nucleus when they are sparse (30-40% confluent) and is retained in the cytoplasm when cells are dense (80-100% confluent). **(B)** Expression of the RGK GTPase REM2 reduces YAP1 nuclear localization while RASL12 does not. Quantification of a transfected cell (marked with 1) and a non-transfected cell (marked with 2) is at right. **(C)** Expression of either RASSF1A or RASSF5A alone does not activate the Hippo pathway, as YAP1 remains predominantly nuclear. Cells transfected with FLAG-RASSFs are marked with a 1 and quantitated at right, a non-transfected control cell is marked with 2. In all panels, scale bar (yellow, Hoechst) represents 10 µm.
